# *dN/dS* dynamics quantify tumour immunogenicity and predict response to immunotherapy

**DOI:** 10.1101/2020.07.21.215038

**Authors:** Luis Zapata, Giulio Caravagna, Marc J Williams, Eszter Lakatos, Khalid AbdulJabbar, Benjamin Werner, Trevor A Graham, Andrea Sottoriva

## Abstract

Immunoediting is a major force during cancer evolution that selects for clones with low immunogenicity (adaptation), or clones with mechanisms of immune evasion (escape). However, quantifying immunogenicity in the cancer genome and how the tumour-immune coevolutionary dynamics impact patient outcomes remain unexplored. Here we show that the ratio of nonsynonymous to synonymous mutations (dN/dS) in the immunopeptidome quantifies tumor immunogenicity and differentiates between adaptation and escape. We analysed 8,543 primary tumors from TCGA and validated immune dN/dS as a measure of selection associated with immune infiltration in immune-adapted tumours. In a cohort of 308 metastatic patients that received immunotherapy, pre-treatment lesions in non-responders showed increased immune selection (dN/dS<1), whereas responders did not and instead harboured a higher proportion of genetic escape mechanisms. Ultimately, these findings highlight the potential of evolutionary genomic measures to predict clinical response to immunotherapy.

## Introduction

Cancer is an evolutionary process, where natural selection acts upon somatic mutations that alter phenotypes, and drives adaptation^1,2^. Recent advances in genomic technologies have enabled the characterisation of mutational landscapes in thousands of malignant^3,4^, and healthy somatic tissues^5,6,7,8^. These studies found that a) 2 to 5 driver mutations are sufficient to initiate a malignancy, b) driver mutations are also present in normal tissue^5,6,7^, c) 90-95%% of somatic point mutations are neutral^7–9^, and d) the signals of negative selection in somatic tissues are weaker compared to germline evolution^7,10^. However, the roles of negative, positive, and neutral evolution during carcinogenesis remains debated^11^, especially with regards to the extent of neutral evolution^12–14^ and negative selection^7,8,15,16^.

The application of evolutionary theory allows us to infer cell growth dynamics, the number of driver alterations^17,18^ and their selective fitness coefficients^19–22^, as well as the impact of deleterious mutations during cancer progression^23,24^. An evolutionary metric recently used to detect selection in cancer studies is the ratio of nonsynonymous to synonymous mutations, *dN/dS*^7,8,25–27^. The rationale is that within a genomic locus, nonsynonymous mutations that decrease cell fitness will show a paucity (negative selection, *dN/dS* < 1) while nonsynonymous mutations that increase cell fitness will be more frequent (positive selection, *dN/dS* >1) compared to synonymous neutral mutations. Mutations modulate fitness by altering the birth-death rate of a cell (driver and deleterious mutations) or by causing immune-mediated predation of the lineage (neoantigens or immunogenic mutations). We recently explored the evolutionary dynamics caused by negative selection operating in cancer, demonstrating that negative selection – and its release by immune escape – leads to a predictable neoantigen variant allele frequency (VAF) distribution^28^. In theory, the shape of the neoantigen VAF distribution can measure selection, but technical limitations around neoantigen detectability in standard genome sequencing make the method impractical and under-powered. Here we show how *dN/dS*-based measures offer a robust means to quantify negative selection strength and detect competing selective forces acting in distinct regions of the cancer genome.

The notion that the immune system influences cancer progression originated in the early 1900s^29,30^. It was only a century later, that studies in mice demonstrated that genetically inbred mice lacking lymphocytes, developed more spontaneous and chemically induced tumors than their wild-type counterparts^30–32^. These results engendered the concept of cancer immunoediting where tumor cells are subject to three phases: elimination, equilibrium, and escape^29^. Cancer immunoediting is an evolutionary process that shapes tumour immunogenicity by selecting for clones depleted of neoantigens (immune-adapted) or with an immune evasion phenotype (immune-escaped)^33–35^. Neoantigens are generated, among other mechanisms, by single nucleotide variants (SNVs) leading to aminoacidic changes in a peptide previously recognized as a self-antigen^36^. However, the extent of immunogenicity derived from SNVs in self-antigens remains unclear, particularly if anchor positions of the wild-type peptide are affected^37^. In our previous work, we observed signals of immune-mediated negative selection in the immunopeptidome, defined as all natively MHC-bound genomic regions, associated to levels of immune infiltration. Nonetheless, a recent study claimed that after applying a more stringent normalization method these regions do not harbour signals of selection^16^. In this work, we corroborated our earlier findings and we further provide an alternative explanation for the lack of signal reported recently.

The recent discovery of immune checkpoints (e.g. PD1 or CTLA4) as mechanism of immune evasion, led to the development of cancer therapies using immune checkpoint inhibitors (ICIs). Despite the promising clinical results of ICIs, only 30% of patients treated with these therapies show significant response. Therefore, considerable effort has been dedicated to understand the interaction between the immune system and cancer^38–43^, and to identify genetic determinants of immunotherapeutic response. To date, quantification of tumor mutation burden (TMB) is the primary genomic biomarker for enrolling patients into ICI treatment. The underlying hypothesis for TMB as a biomarker is that a higher number of somatic mutations leads to a higher number of neoantigens, and therefore a higher likelihood of immune clearance after checkpoint inhibition. However, recent studies have shown that even mismatch repair proficient tumors display a pathological response^44^, emphasizing the need for quantifying the true immunogenicity of the cancer genome and their potential clinical response to immunotherapy.

Here, we modelled cancer initiation and progression by adapting a stochastic branching process^45^ to simulate changes in *dN/dS* over time as a measure of selection and tumor immunogenicity during immunoediting. Using the insight gained from our model, we assessed *dN/dS* values in 8543 primary tumours, as well as 308 metastatic cancers treated with ICIs. We first corroborate that immune *dN/dS* correlates with levels of tumor infiltrating lymphocytes – a measure of the strength of immunoediting - in non-escaped tumors. Finally, by estimating immune *dN/dS* in pre-treated patients, we reported clinical response in immune-escaped patients that had an absence of immune selection (immune *dN/dS* ~ 1). In contrast, tumors with low immune *dN/dS*, and therefore low levels of tumor immunogenicity, did not respond to the action of immune checkpoint inhibitors.

## Results

### A mathematical model of immunoediting

We extended our previous modelling work to incorporate the acquisition of nonsynonymous and synonymous mutations in driver (positively selected) and passenger (neutral) loci^24,46,47^, as well as in regions exposed to the immune system and regions that confer immune-evasion properties (Fig 1). The interaction of different mutations and the observed evolutionary dynamics can be simplified into four phases: 1) A pre-neoplastic phase where cells do not have cancer driver mutations but may acquire passenger, immunogenic or escape mutations (Fig 1A), 2) a neoplastic phase that begins when a driver mutation avoids stochastic drift and initiate a clonal expansion (Fig 1B), 3) an elimination phase where cells acquiring somatic mutations recognized by the immune system are eliminated (Fig 1C), and 4) a phase where expanding clones lead to a clinically-detectable tumor through either depletion of immunogenic mutations (immune adapted) or through a mutation in the genome that triggers an immune escape mechanism (immune escaped) (Fig 1D). An activated escape mechanism hides the clone from the immune system so that neoantigens accrue without being depleted by negative selection raising the overall tumor immunogenicity (Fig S1A). It is possible for these phases to overlap each other. For example, an escape mutation occurring pre-driver acquisition and thus pre-clonal expansion leads to tumors “born” immune-escaped.

**Figure 1.**
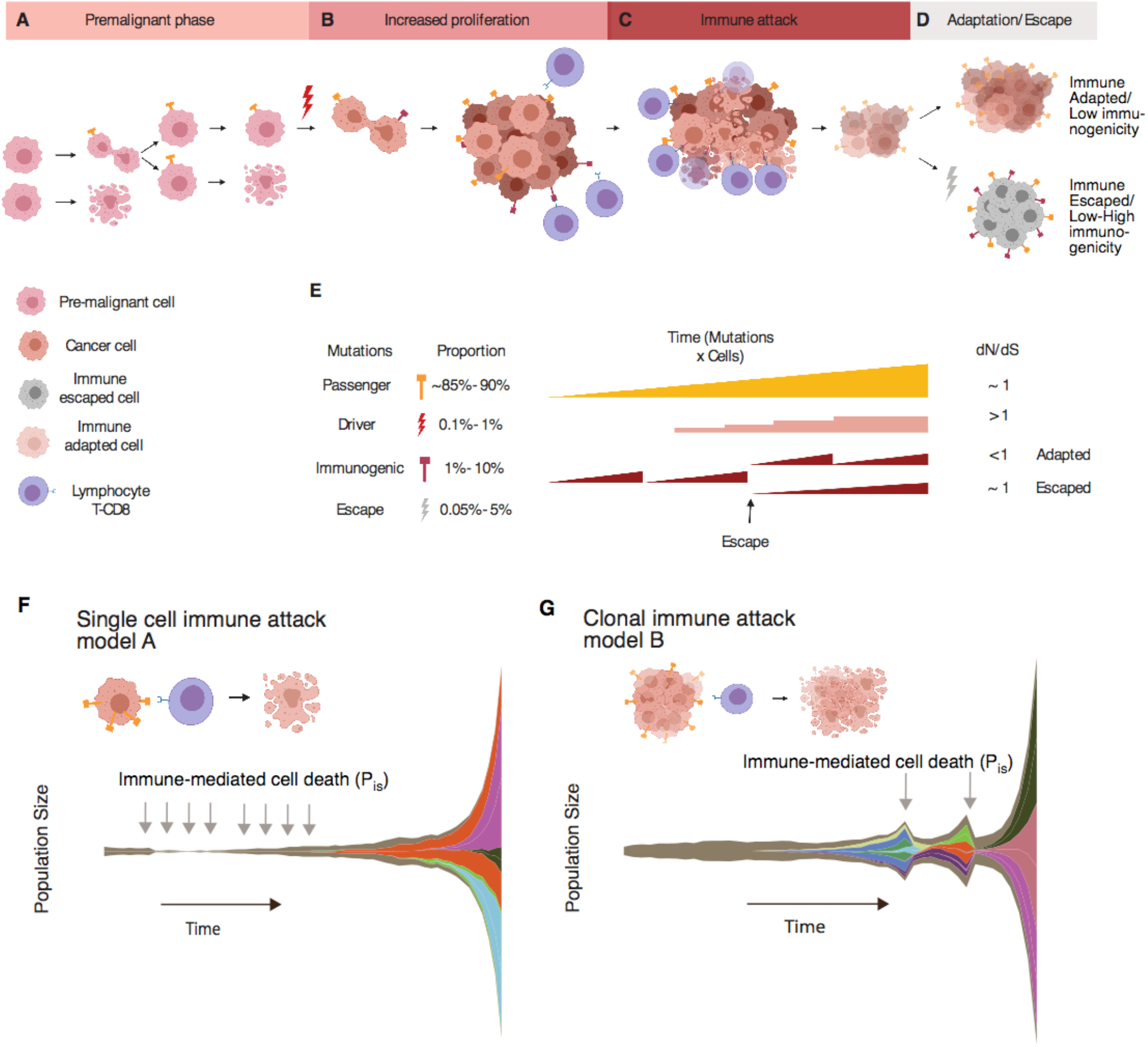
Description of the stochastic branching process used to model immunoediting. A) An initial set of wild type cells (Pre-malignant cell) divide and accumulates mutations. B) A driver mutation increases the probability of cell division initiating a phase of increased proliferation of clones (Cancer cell). C) During the phase of immune attack, the immune system removes cells carrying immunogenic mutations and might eradicate the tumor completely or force the tumor to adapt or escape. D) Two possible scenarios emerge as the outcome of immunoediting, cancer cells survive not harbouring immunogenic mutations (Immune adapted) or due to the acquisition of an immune evasion mechanism (Immune escaped). E) These scenarios can be differentiated by looking at the ratio of nonsynonymous to synonymous mutations (dN/dS) in immune exposed regions of the genome. We defined two hypotheses of immune recognition: F) Single cell immune attack where any single cell carrying a neoantigen is able to initiate an immune response and be eliminated at a rate of immune-mediated cell death of P_IS_, G) Clonal immune attack where a minimum percentage of immunogenic cells is needed to elicit an immune response, as recently observed in mice models ^48^.

We initiated our model in the pre-neoplastic phase with a pool of *N* cells having an equal probability of birth (*b*) and death (*d*): *b=d=0.5* (Methods). For each successful cell division, a number of new mutations are sampled from a Poisson distribution with mean μ×L (mutation rate measured in mutations per base pair per cell division multiplied by the length of the coding genome, L). We introduced nonsynonymous and synonymous mutations at a constant relative rate of 3 to 1 given the expected genome composition^49^, so we could calculate the ratio between these two types of mutations (*dN/dS*) in the evolved population of tumour cells. We assumed that passenger nonsynonymous and all synonymous mutations are neutral. Once a cell acquired a nonsynonymous mutation in a driver, the probability of cell division *b* increases by a fixed value obtained from a Gompertz function (Methods), driving the next stage of tumorigenesis.

During immunoediting^29,50^, cells carrying an immunogenic mutations may elicit an immune response. We tested whether or not *dN/dS* values derived from the immunopeptidome, the portion of the genome constantly exposed to immune recognition (defined as ‘Immune *dN/dS*’), quantifies overall tumor immunogenicity, and differentiates between adaptation and escape. We expected that when the immune predation was active and there were no escape mechanisms evolved, the immune *dN/dS* would be lower than 1 showing overall low tumor immunogenicity. Conversely, in the presence of escape mechanisms immune *dN/dS* would have values closer to 1, and therefore high immunogenicity. Additionally, we could also measure a ‘global *dN/dS*’ by using mutations in all loci of the genome, and a ‘driver *dN/dS*’ by considering only mutations in driver loci (Fig 1E). We then modelled two hypotheses of immune recognition (Fig S1B): (1) a classic model (model A) where a single cell carrying an immunogenic mutation is sufficient to elicit an immune response (Fig. 1F), and (2) a clonal model (model B), recently suggested^48^, where a percentage *Pclonesize* of the total cells carrying the same immunogenic mutation is needed for the immune system to attack (Fig. 1G). In model A, the immune system is constantly pruning immunogenic cells, whereas model B produces a “rise and fall” pattern where immunogenic cells are allowed to expand to a threshold size but are then eliminated, similar to mass extinction events. Cells bearing a neoantigen are killed at an immune-mediated cell death rate P_IS_, where P_IS_ ∈ [0,1]. This parameter models the stochastic probability of encounters between antigen presenting cells and cytotoxic T-cells. Model parameters are summarized in Supplementary Table 1.

### Evolutionary dynamics of *dN/dS* during immunoediting reveals genomic signals of tumor immunogenicity

To first understand *dN/dS* dynamics during the pre-neoplastic phase, we simulated the acquisition of neutral mutations only (non-synonymous passenger and synonymous mutations) in an initial population of 32 cells for 30 generations (Fig S2). We compared three mutation rate regimes similar to those founds in some neoplasms: microsatellite stable (μMSS*=10^−8^* mutations/bp/division), microsatellite unstable (μMSI*=10^−7^*), and POLE-like (POLE*=10^−6^*). On average, the population size remained constant over time for the three regimes and the number of mutations was higher for higher mutation rate regimes (Fig S2A-B). The average number of mutations per simulated population was 10^2^, 10^3^, and 10^4^ for each mutation rate regime respectively (Fig S2B). As expected under neutral dynamics, we observed that the average *dN/dS* did not deviate significantly from 1 and the variance was lower at high mutation rates. (95%CI for *10^−8^*: 0.54-2.31, MSI:0.79-1.30, POLE:0.91-1.06) (Fig S2C).

To determine the influence of positive selection on *dN/dS* values over time during the increased proliferation phase, we simulated only passenger and driver events. We simulated 1000 datasets assuming 0.1%, 0.5% and 1% of driver sites (Fig S3). We focused our analysis on simulations where a clonal expansion occurred, as defined by a growing population of more than 1000 cells within 100 generations (Fig 2A). We calculated *dN/dS* over time for all mutations (global *dN/dS*) and for only driver mutations (driver *dN/dS*). We observed large fluctuations of the global *dN/dS* values among the first generations due to the low number of mutations (Fig S3D). Interestingly, the accumulation of neutral variants pushed global *dN/dS* values to 1. Driver *dN/dS* peaked at high values and subsequently decreased towards one due to the accumulation of low frequency neutral variants (Fig S3E). As we demonstrated in Williams et al^21^, mutation frequency and driver *dN/dS* are expected to be positively associated showing the highest values at the largest clone sizes (Fig 2B). Accordingly, and as observed recently in clonal hematopoiesis^22^, the allele frequency spectrum (Cancer Cell fraction or CCF) of synonymous and non-synonymous mutations (Fig. 2C) showed that the observed high driver *dN/dS* is a consequence of proportionally fewer synonymous mutations at higher CCF thresholds compared to nonsynonymous mutations.

**Figure 2.**
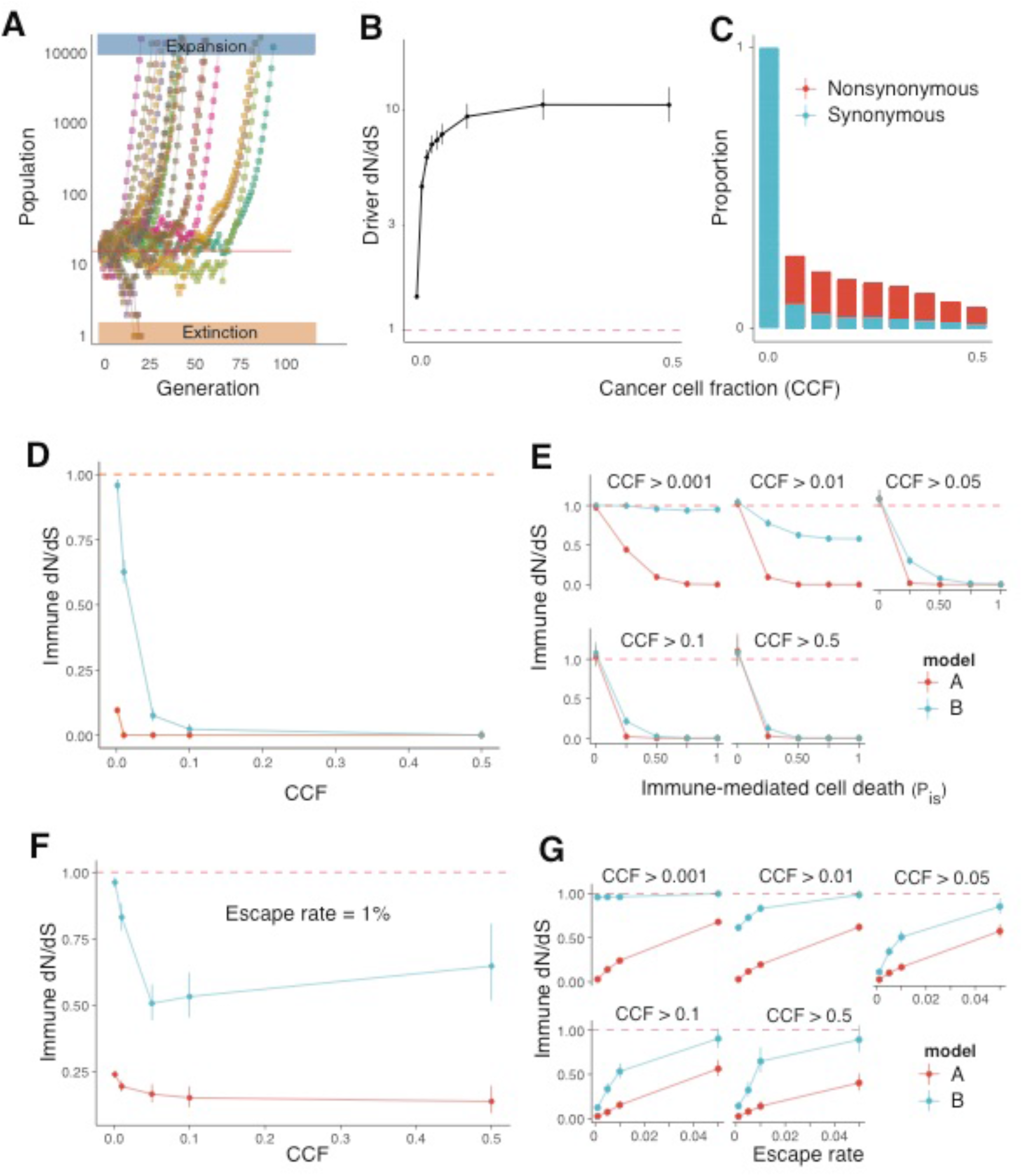
Immunoediting leads to tumor adaptation or escape. A) We defined two outcomes for each simulation: expansion and extinction. Expansion: Clonally expanded populations (Blue) that reached an upper limit of number of cells in the first n generations. Extinction: Simulations that drifted to extinction among the first n generations (Orange). B) driver dN/dS relationship to the cancer cell fraction. As described in Williams et al^21^ we show in our model that driver dN/dS increases at increasing values of clonality. C) Relative proportion of nonsynonymous to synonymous mutations. The upward trend of dN/dS is due to a high proportion of synonymous mutations removed at increasing CCF cut-offs. D). Immune dN/dS relationship to cancer cell fraction for single cell (model A, Red) and for the clonal model (model B, Blue). A sharp decrease in dN/dS at increasing CCF cut-offs consistent with the theoretical predictions for strong negative selection^21^. E) Immune dN/dS relationship to the probability of immune-mediated cell death at different levels of CCF. At low CCF, the dN/dS for model B is closer to one across all levels of immune death due to the presence of several undetected small frequency clones carrying neoantigens. At high CCF, both models show strong association between immune death and dN/dS. This results into cancer clones depleted of neoantigens, classified as immune-adapted and bearing an overall low tumor immunogenicity. F) Immune dN/dS at different CCF cut-offs when including escape mutations at 1% rate. At low CCF levels, immune dN/dS decreases when increasing CCF but escaped clones push the signal of immune dN/dS towards one at high CCF cut-offs for model B. G) Immune dN/dS relationship to the probability of immune-mediated cell death at different levels of CCF when escape mutations are included. For both models, increasing the probability of escape events pushes dN/dS values back to one for all CCF cut-offs, reflecting a relaxation of immune-mediated negative selection. Ultimately, these tumors are growing with escape mechanisms that allow the accumulation of neoantigens that increase the overall tumor immunogenicity.

During the elimination phase, in addition to driver and passenger mutations, we introduced immunogenic mutations (5% of immunogenic sites) and explored the dynamics under two mechanisms of immune recognition (Single cell versus clonal immune attack). We first calculated immune *dN/dS* values at different cancer cell fraction (CCF) cutoffs. We observed that at increasing clone sizes the immune *dN/dS*, and therefore tumor immunogenicity, value was approaching zero for both models (Fig 2D). As in model B negative selection is absent for small clones (low CCF), immune *dN/dS* was closer to 1. Then, we calculated immune *dN/dS* at varying rates of immune-mediated cell death, P_IS_, for different clone sizes (Fig 2E). We first confirmed that when the immune system was inactive (P_IS_ = 0), the immune *dN/dS* was one for all clones. At increasing levels of effective immune surveillance both models demonstrated depletion of immunogenic mutations, and therefore low levels of tumor immunogenicity. Immune *dN/dS* in model B was less affected by this parameter given that multiple immunogenic mutations can remain hidden at low frequency. Ultimately, these simulations showed how immune *dN/dS* reveals the action of immune-mediated negative selection and can be used as a proxy for tumor immunogenicity.

We next explored immune *dN/dS* values during the evolution of immune escape. The activation of escape is modelled as a stochastic event occurring at a fixed rate that depends on the proportion of escape sites in the genome. We repeated simulations using an immune-mediated cell death of P_IS_ = 1 at different rates of escape. We first found that when the proportion of escape sites was 1%, immune *dN/dS* captured the action of immune-mediated negative selection across the whole frequency spectrum (Fig 2F). In model B, immune escape pushed immune *dN/dS* values back to one, slightly increasing overall tumor immunogenicity. At higher rates of immune escape, we observed increased immune *dN/dS* demonstrating how tumor immunogenicity is restored for all clone sizes when escape events are more common (Fig 2G). Notably, when the escape rate was 5%, all clone sizes in model B reached immune *dN/dS* values close to one, highlighting high levels of tumor immunogenicity. By acquiring escape mechanisms, negative selection in the immunopeptidome is relaxed, the accumulation of immunogenic mutations becomes neutral, and tumor immunogenicity is restored.

The results of our modelling provide a theoretical framework of co-evolution of somatic cells and the immune system and a basis to quantify tumor immunogenicity based on immune *dN/dS*. Further, it illustrates the importance of choosing an appropriate region of the genome to analyse immune selection and how clone sizes explain different levels of tumor immunogenicity. Moreover, we speculate that when mixing patients that are immune-escaped with non-escaped, signals of immune-mediated negative selection are no longer representative of the overall tumor immunogenicity.

### High levels of lymphocyte infiltration are associated to strong immune-mediated negative selection and low levels of tumor immunogenicity

To measure global, driver, and immune *dN/dS* values using genomic data, we developed SOPRANO (Selection On PRotein ANOtated regions), a bioinformatic pipeline that measures the extent of selection in specific regions of the genome (github.com/luisgls/SOPRANO). It extends our previous work, where we calculated *dN/dS* corrected for mutational context using a 7-substitution type (SSB7) or a 192-substitution model (SSB192)^8^. Here, we have extended the method to account for any set of concatenated genomic regions allowing for patient- and region- specific *dN/dS* estimates. We applied SOPRANO to 8543 tumour samples from 19 cancer types from The Cancer Genome Atlas (TCGA), using the SSB192 model (Fig 3, Supplementary Table 2). We compared the ratio of *dN/dS* values between regions inside and outside the immunopeptidome (ON/OFF *dN/dS*). We defined the immunopeptidome as all possible wild-type 9-mer regions present in the genome of a patient that are predicted to bind to the MHC-I complex with an affinity of %rank < 0.5 as defined in netMHC4.0 (Fig S4). In our first analysis, we used our previously published set of regions that bind to HLA-A0201 and compared them to a recently published proto-HLA^16^ consisting of multiple HLA alleles. We found that lung adenocarcinoma (LUAD) and melanoma (SKCM) showed a depletion of nonsynonymous mutations in HLA-A0201 binding regions, and that LUAD, HSNC and LUSC showed a depletion of nonsynonymous mutations in proto-HLA regions (Fig 3A). We compared the immune *dN/dS* values (ON/OFF *dN/dS*) obtained using SOPRANO to the values of immune selection (normalized HLA-binding mutation ratio or HBMR), recently reported by Van Den Eynden et al^16^. We observed a significant correlation between the ON/OFF *dN/dS* ratio and the reported normalized HBMR using the proto-HLA (R=0.77, P Value= 0.00054, Fig 3B) but not when comparing to the HLA-A0201 (R=0.37, P Value= 0.15, Fig S5). Expectedly, the correlation for the HLA-A0201 was lower given the HBMR value was calculated using multiple HLAs- binding regions and therefore every patient not carrying the proto-HLA allele will contribute with only neutrally accumulating mutations. In consequence, it is important to note that the smaller the fraction of the assessed region that is truly under immune selection, the more neutral the *dN/dS* value would appear.

**Figure 3.**
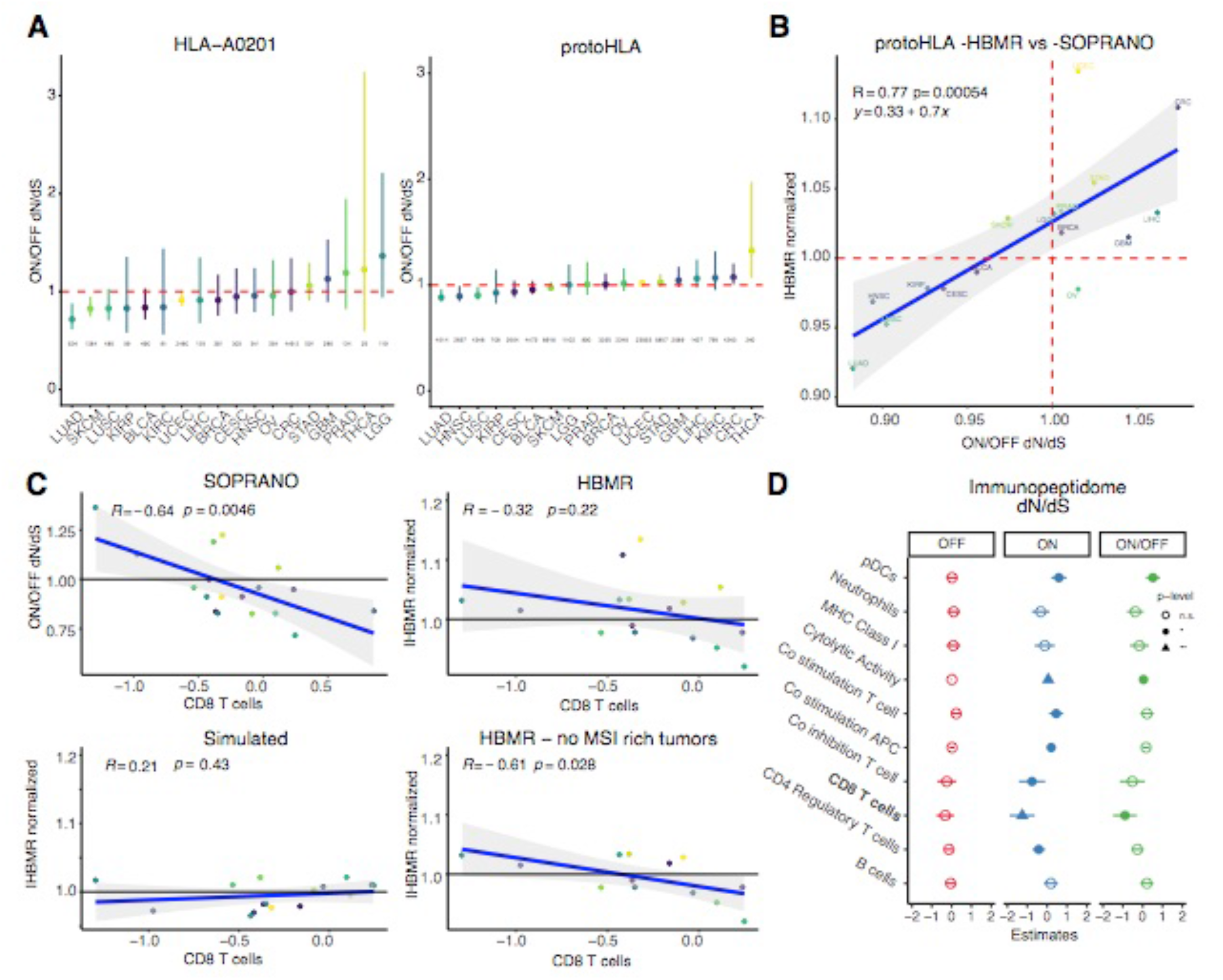
Immune dN/dS and immune activity across multiple tumor types. A) Immune dN/dS (ON/OFF dN/dS ratio) in multiple tumor types using either a curated HLA-A0201 target region or a proto-HLA consisting of the most common HLA haplotypes in the population obtained from Van Den Eynden et al^16^. Numbers represent the mutations ON target for each dataset. B) Comparison of immune dN/dS values using SOPRANO SSB-192 and normalized HBMR values reported in Van Den Eynden et al.^16^ C) Linear regression models for immune dN/dS and HBMR values versus median CD 8 T cell infiltration levels. In the analysis with no MSI-rich tumors, in addition to colorectal (CRC), we removed Stomach and Uterine cancer (STAD and UCEC). D) Linear mixed model using dN/dS values as the dependent variables and all immune metrics as independent variables. Model selection using AIC revealed that ON/OFF dN/dS is strongly associated to the levels of CD8 T cells. No immune value was associated to the global dN/dS (OFF).

To determine whether immune-mediated negative selection was associated with levels of immune activity, we compared immune *dN/dS* to the levels of immune infiltration previously reported in TCGA data^51^ (Fig 3C). Median CD8 T cells significantly correlated to the SOPRANO-derived immune *dN/dS* values in HLA-A0201 regions (p=0.0046) but not to the HBMR values (proto-HLA) calculated in ^16^ (p=0.22), even though the trend was negative for both. As expected, the correlation was also not observed in the simulated dataset. Interestingly, when tumour types where microsatellite instability (MSI) and mismatch-repair deficiency was common, such as colorectal (CRC), stomach (STAD) and uterine cancer (UCEC), were excluded from the analysis, the correlation between proto-HLA HBMR and the median CD8 T cells was significant (P=0.028), indicating that negative selection acts differently in these different tumour subgroups. This makes sense as hypermutant MSI tumours have a large frequency of escape events, such as upregulation of immune checkpoint mechanisms, loss of heterozygosity in the HLA region or mutations in genes associated to the antigen presenting machinery^28,33,52^. This last correlation was also strongly significant for cytolytic activity (P-value = 6e-04, Fig S6).

We applied a linear mixed model to determine the contribution to the global *dN/dS* (OFF), the immunopeptidome-specific (ON) and the immune-*dN/dS* (ON/OFF) using reported immune variables (Fig 3D). We performed a stepwise model selection, and the initial (Fig S7) and best performing model for predicting Immune *dN/dS* (R-square adj= 0.89, AIC= −83, p-value = 0.01) had CD8 T cells as the most significant explanatory variable. Importantly, none of the variables could explain global *dN/dS* values and seven out of the ten variables tested was significantly associated to the immunopeptidome-specific ON value. Moreover, we found that there was no significant correlation between CD8 T cells and immune *dN/dS* in patients that have a truncating mutation in a gene associated to the antigen presenting machinery or genes defined as escape genes previously^34^.

In summary, these results highlight the importance of considering multiple confounding factors when drawing conclusions about the absence of negative selection at the cohort level using *dN/dS*. These results further suggest that high mutation burden tumors show signals of relaxed immune-selection confounding the calculation and interpretation of *dN/dS* probably due to the presence of acquired escape mechanisms, as our theoretical model predicts.

### Immune-escaped tumors show relaxed immune-mediated negative selection and high tumor immunogenicity

Following our immunoediting model, we hypothesized that escape events mask the signal of immune mediated negative selection and restore tumor immunogenicity. We ran SOPRANO using a patient specific immunopeptidome (private HLA alleles) in colorectal (CRC), stomach (STAD) and uterine cancers (UCEC). While tumor mutation burden was expectedly higher for MSI and POLE tumors^53^ (Fig S8), ON-target immunopeptidome *dN/dS* values for MSI and POLE subtypes were also higher than for MSS tumors (Fig S9), consistent with high-mutation rate tumours being very frequently immune-escaped^28^.

We then classified different escape mechanisms for these tumors based on previous work^28^. We found that immune-escaped (Escape+) tumors have significantly more somatic mutations compared to non-escaped (immune adapted) tumors (MSS p=0.0085 and MSI p=0.00026, Fig 4A). We then reasoned that a larger number of mutations in MSS escape+ tumors would push immune *dN/dS* towards 1, given an extended time of neutral mutations accumulating in the genome *after* immune escape has occurred. Indeed, we found a significant positive correlation between tumor mutation burden and immune *dN/dS* for MSS escape+ tumors but not for immune adapted tumors (Escape-, p=1e-04, Fig 4B), suggesting that immune selection was still active in patients without an escape mechanism. We observed the same results when restricting our analysis to only clonal mutations (P Value = 0.003, Fig S10), confirming our previous suggestion that immune-escape tends to occur early in the genesis of these malignancies^28^.

**Figure 4.**
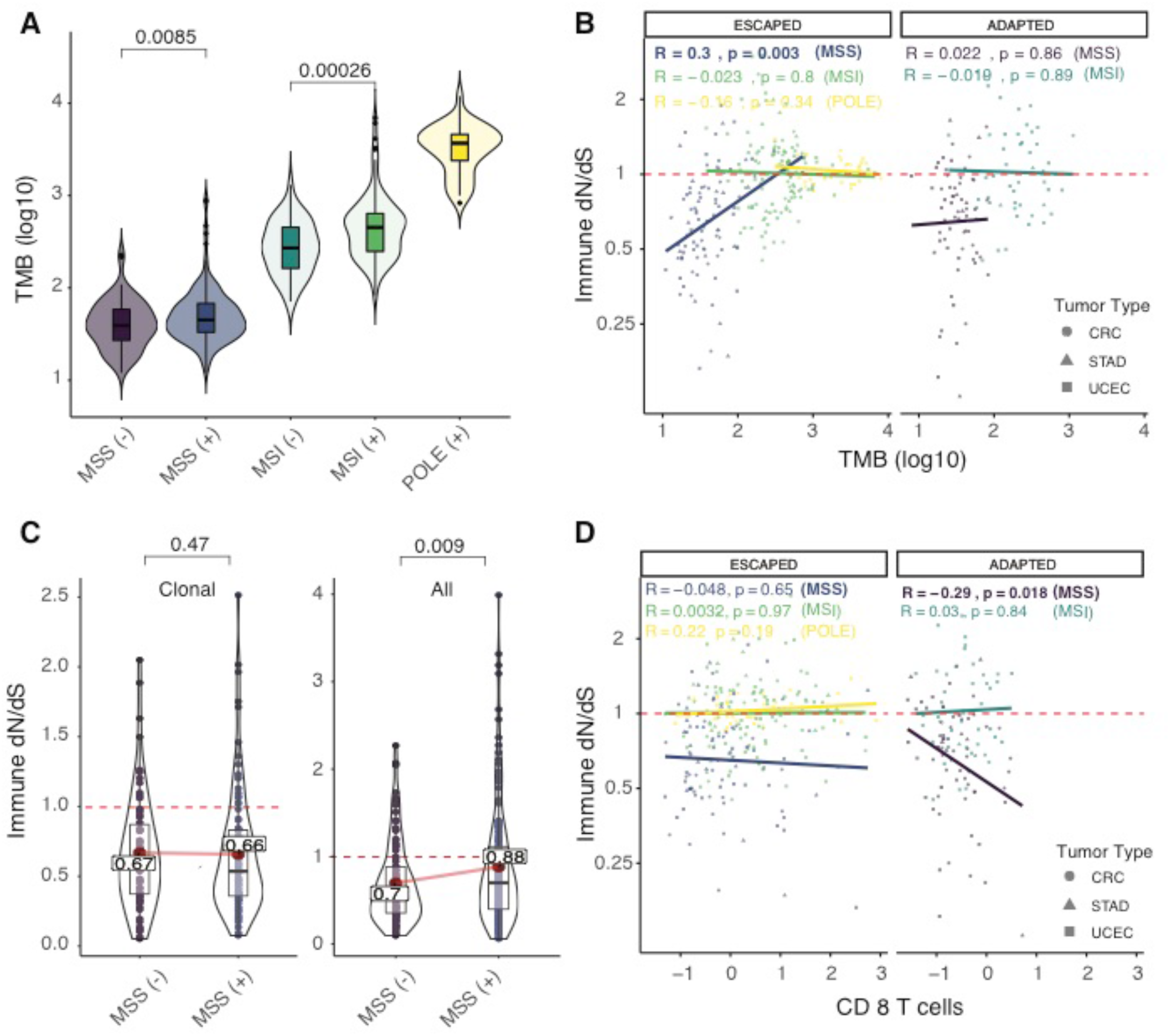
Patient specific analysis of colorectal (CRC), stomach (STAD) and uterine cancer (UCEC) using immune dN/dS. A) Tumor mutation burden (TMB) for different subtypes of cancers, including Microsatellite Stable (MSS), Microsatellite Instable (MSI) and POLE mutants, classified as immune-adapted (−) or immune-escaped (+) based on the presence of escape mechanisms (obtained from Lakatos et al^28^). B) Relationship between Immune dN/dS values to TMB, following the same classification for patients as in A. C) Comparison between immune dN/dS values for immune-escaped and immune-adapted MSS tumors using all or only clonal mutations. D) Relationship between immune dN/dS values and the reported CD 8 T cell infiltration following the same classification for patients as in A and B.

Our theoretical model predicted that immune *dN/dS* will remain lower than one when at large clone sizes in non-escaped patients. We expected that clonal mutations may still hold the signature of negative selection (active before escape) while subclonal mutations would be freely accumulating in immune-escaped tumors. Consequently, when we compared immune *dN/dS* between immune-adapted (escape-) and immune-escaped (escape+) MSS tumors, we observed that immune-escaped tumors had immune *dN/dS* values significantly closer to 1 compared to immune-adapted tumors when using all mutations, but not when using clonal mutations (0.88 versus 0.7, Wilcoxon signed rank test=0.0009, Fig 4C). In the case of immune-adapted tumors, the Immune *dN/dS* when using all or clonal mutations remained similar (0.68 versus 0.70 immune *dN/dS*) while immune-escaped tumors had a significantly higher immune *dN/dS* when using all mutations (0.88 versus 0.66 immune *dN/dS*, P Value=0.007) (Fig S11).

To validate that the strength of immune-mediated negative selection depends on immune activity, we compared the patient specific immune *dN/dS* to the CD8 T cell infiltration (Fig 4D). We found a significant association between immune activity and CD8 T cells in immune-adapted MSS tumors (P Value = 0.018), reaffirming that native HLA binding regions hold information on the strength of immune selection elicited by CD 8 T cells. Interestingly, MSI tumors without an annotated escape mechanism did not follow this pattern suggesting that these tumors may have an unknown escape mechanism. These results highlight the importance of understanding the evolutionary dynamics of tumors under immunoediting and provide a theoretical explanation of why tumors with high mutation burden are better candidates for immunotherapy. Such tumors have an overall higher tumor immunogenicity that can be quantified using *dN/dS* in the immunopeptidome.

### Immune-escaped tumors have better response to immunotherapy than immune-adapted tumors

To finally address the clinical importance of escape mutations and immune *dN/dS* as a surrogate of tumor immunogenicity, we analysed 308 metastatic cases subjected to immunotherapy with checkpoint inhibitors mainly with Ipilimumab, Nivolumab, Ipi+Nivo, and Pembrolizumab from the Hartwig Medical Foundation cohort^4^ (Fig 5A). The specimens were sequenced before treatment was started. Following RECIST guidelines patients were classified into complete and partial response and into progressive or stable disease (Methods). There were 78 responders recorded (Partial or complete response) and 229 non-responders (Progressive or Stable disease). Due to the unavailability of patient specific HLA, we calculated immune *dN/dS* using HLA-A0201 and observed a lower immune *dN/dS* for non-responders compared to responders, suggesting that patients with no response to immunotherapy were already adapted to the action of immune response (Fig 5B). Next, we assembled a list of escape genes associated to the immune response and further classified patients into immune-escaped and non-escaped (Methods). Given that only genomic data was available for this cohort, we could only classify patients into genetic escape and not into other immune evasion events, such as overexpression of immune checkpoint inhibitors. We found that the proportion of responders with a genetic escape mechanism was significantly higher compared to non-responders (Chi-square P value = 0.001, Fig 5C), indicating that escape mechanisms were independently associated to the clinical response during immune checkpoint therapy.

**Figure 5.**
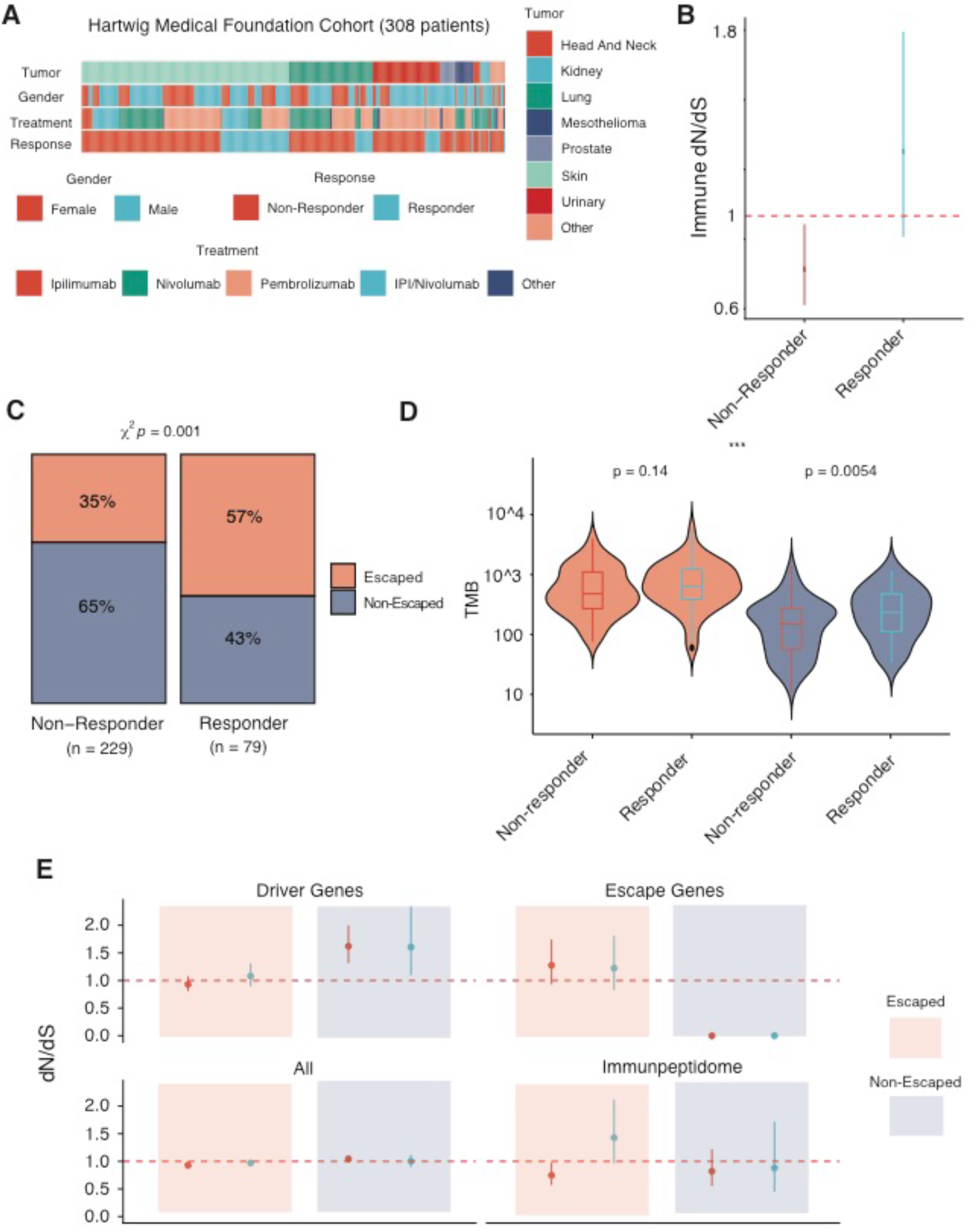
Analysis of Hartwig Medical Foundation metastatic cohort under immunotherapy. A) 308 patients with information about their response post-immunotherapy were selected. B) Immune dN/dS values for responders and non-responders reveals immune dN/dS lower than one for non-responders consistent with an overall tumor immunogenicity unresponsive to immunotherapies. C) Proportion of escaped and non-escaped tumors classified by clinical response. Responders are enriched in genetic escape mechanisms. D) Tumor mutation burden (TMB) for escaped and non-escaped tumors classified by response status. Among non-escaped patients, responders have a higher TMB than non-responders. E) dN/dS values for driver genes (196 genes from Martincorena et al ^7^), escape genes, all the exome and for the immunopeptidome. The panel show positive selection in driver genes for escaped tumors but not for non-escaped tumors. dN/dS in escape genes was indicative of positive selection for escaped tumors. Non-escaped tumors were selected for not having non-synonymous mutations in escape genes, hence dN/dS was 0. Global dN/dS values were one for all cases consistent with the lack of selection in the majority of the coding genome. dN/dS in the immunopeptidome was indicative of immune-selection for all groups except for escaped tumors in responders.

Given that tumor mutation burden (TMB) is the current FDA-approved prognostic marker of immunotherapy, we compared TMB between responders and non-responders. As expected, we found that responders had a significantly higher TMB than non-responders before the treatment (Fig S12A, P-value = 9.8*e-6, U Mann Whitney). In parallel, we looked at TMB between escape and non-escaped patients and found that escaped patients had also a significantly higher TMB compared to non-escaped (Fig S12B, P-Value < 2.2e-16, U Mann Whitney). We also explored if TMB was different within escaped and non-escaped groups separated by response. We found that TMB was significantly higher for responders among the non-escaped group (P Value=0.0054, U Mann Whitney) but not different among escaped patients (P value=0.14, U Mann Whitney) (Fig 5D). The fact that, among non-escaped patients, responders had higher TMB, suggest that a group of responders had an escape mechanism that was not considered in our classification. This is expected given that we did not consider all possible escape mechanisms such loss of HLA heterozigosity^33^, epigenetic escape such as transcriptional silencing by changes in methylation^34^, or extrinsic factors such as the accumulation of dysfunctional T cells^54^, all mechanisms of immune evasion recently described in the literature.

Finally, we calculated *dN/dS* for driver, global, escape, and immune regions in these four groups (Fig 5E). We found that the driver *dN/dS* was positive for non-escaped tumors as expectedly, but surprisingly neutral for immune-escaped tumors. The escape *dN/dS* showed signals of positive selection for escaped patients and given that no nonsynonymous escape mutations were present in non-escaped patients, the escape *dN/dS* was zero. The global *dN/dS* was consistently close to one for all groups. Importantly, among escaped patients, while the TMB was not different between responders and non-responders (Fig 5D), the immune *dN/dS* of non-responders was lower than one and lower than the immune *dN/dS* of responders. Ultimately, this validates immune-adaptation in non-responders showing less neoantigens and therefore low levels of tumour immunogenicity for immunotherapies to have an effect.

Overall, our results highlight the importance of properly stratifying patients based on escape mechanisms and immune *dN/dS* for a correct interpretation of the evolutionary dynamics of tumors. In the future, these genomic based classification in combination with current standard practices, could be used as prognostic biomarkers for checkpoint inhibitor immunotherapies^55^.

## Discussion

The remarkable clinical response demonstrated by immune checkpoint inhibitors (ICIs) has led to a growing interest in understanding the interactions between cancer and immune cells^33,35,56–58^. Although immunoediting is widely recognized as an evolutionary process that selects for clones with low immunogenicity or clones with an escape mechanism, its dynamics in the context of carcinogenesis and response to treatment are poorly understood. During immunoediting, growing cells are subjected to immune-mediated negative selection, shaping the landscape of mutations observed in cancer. However, negative selection in cancer has been a controversial topic^10,15,16^. While some studies have shown evidence of an association between immune activity and selective pressures^8,34,51,56,59^, others have claimed that there is a lack of evidence to prove this relationship^16^. Given that several studies have applied *dN/dS* as a metric of selection in cancer and in normal tissue^7,8,60–63^, we aimed to prove the use of *dN/dS* in the immunopeptidome as a proxy of tumor immunogenicity and as a potential biomarker of immunotherapeutic response. In brief, we show that immune *dN/dS* quantifies the extent of negative selection exerted by the immune system and how levels of tumor immunogenicity measured by immune *dN/dS* can be used as a genomic biomarker for response to immunotherapy.

We first show the evolutionary dynamics of tumorigenesis under two radically different outcomes of immunoediting, immune-adaptation and immune-escape. Such distinction is a key feature of cancer evolution and has profound clinical implications. Immune-adapted tumors can only emerge in tissues where the immune system can exert a selective pressure, suggesting that tissues with a high capacity of immune recognition (immune-competent) are more likely to generate clones with a depletion of immunogenic mutations if the probability of escape is low (i.e. a low mutation rate). A lower number of neoantigens could allow tumors to grow in both low and high immunogenic tissues, potentially making them more aggressive when colonizing new niches. Supporting our hypothesis, a recent study of longitudinal recurrence of metastasis reported a more aggressive phenotype in metastatic deposits that had higher levels of immune-selection^56^. However, whether tumor cells growing in immune-competent tissues are more likely to colonize new niches and how long it takes those tumor cells to readapt or to find a novel escape mechanism, as has been previously observed in mice models^29,31,50^, remains a challenging question.

Our immunoediting model predicts that immune-adapted tumours have an overall low tumor immunogenicity and will be less likely to respond to ICIs regardless of tumor mutation burden status (TMB). TMB has been regarded as a measure of tumor immunogenicity and is the current FDA-approved prognostic biomarker used to enrol patients for ICI treatment. However, TMB does not capture the full evolutionary history of the tumor and several patients do not respond despite their TMB status. In addition, a recent study has shown that mismatch repair-proficient colorectal cancers can also achieve clinical response^44^. Motivated by this, we propose that immune *dN/dS* can be used, in addition to TMB and escape mechanisms, to stratify patients into adapted and escaped. As evidence of this, we demonstrate that in a metastatic cohort, non-responders have an immune *dN/dS* lower than one prior to immunotherapy and are thus immune-adapted, whereas responders have immune *dN/dS* values of one, and are more likely to be immune-escaped.

In conclusion, our study reflects the importance of understanding the evolutionary dynamics of immunoediting during tumor evolution and how immune selection edits the genome of tumor cells. Differentiating between immune adapted and immune escaped tumors is a key factor when predicting which patients will benefit from immunotherapies. In the future, we believe that immune *dN/dS* can be used as read-out of tumor immunogenicity, that, in combination with other prognostic measurements, can be used to predict response to immunotherapy.

## Acknowledgements

L.Z. is supported by the European Union’s Horizon 2020 research and innovation programme under the Marie Skłodowska-Curie Research Fellowship scheme (846614). A.S. is supported by the Wellcome Trust (202778/B/16/Z) and Cancer Research UK (A22909). We acknowledge funding from the National Institute of Health (NCI U54 CA217376) to A.S. and T.G. This work was also supported by a Wellcome Trust award to the Centre for Evolution and Cancer (105104/Z/14/Z). We thank Claire-Alix Garin and Marco Punta for support and discussion. This publication and the underlying study have been made possible partly on the basis of the data that Hartwig Medical Foundation and the Center of Personalised Cancer Treatment (CPCT) have made available to the study.

## Author contributions

LZ conceived, designed, implemented and performed all analysis, GC supported with the model implementation, MW, EL, and BW provided support with the mathematical inferences of the model. KAJ provided bioinformatic support. TG and AS supervised the project. LZ wrote the first draft of the manuscript. LZ, AS and TG wrote the final version of the manuscript with the help of all the authors.

## Data availability

TCGA data was obtained from GDC portal and processed as described in^13^. Assembled list of escape mechanisms for COAD, READ and STAD and UCEC was obtained from Lakatos et al^28^. Hartwig Medical Foundation data was downloaded from Hartwig Data Portal. HBMR values of selection in the immunopeptidome were obtained from the supplementary material in Van Den Eynden et al^16^. Normalized scores for immune cell infiltration was obtained from Rooney et al^51^.

## Code Availability

SOPRANO is freely available at github.com/luisgls/SOPRANO. Simulator of stochastic branching process for immunoediting is available at github.com/luisgls/dNdSSimulator upon request. All figures are available as a markdown file.

## Methods

### 1.0 Evolutionary model of tumorigenesis under immunoediting

#### 1.1 Computational model

We have developed a discrete-time non-spatial stochastic branching process of somatic evolution. It models the acquisition of somatic mutations and their associated effect on the phenotype of single cells. The model can be initialized with any number of wild-type single cells and a set of initial parameters described in supplementary table 1A.

#### 1.2 Cell division process

The model simulates cellular proliferation starting from *x*_2_ identical initial cells available at time *t*_2_ = 0, that divide synchronously; each time-step therefore is represented in units of tumour doublings or generations, as in earlier works^47^.

At every time-step every single cell (SC) in the model undergoes a stochastic process with a probability that depends on a parameter *p*_6_. The outcomes of this process are either zero, one, or two single cells in the model:

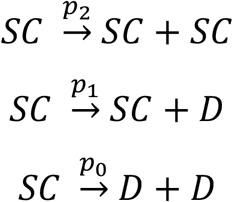

where *p*_2_ + *p*_@_ + *p*_A_ = 1 and D denotes “dead” cells. We consider death as any process that removes the cell from the dividing population, such as apoptosis, senescence, quiescence, or differentiation. To simplify the possible outcomes of the model, we consider *p*_@_ = 0. Thus, our branching process consists only of no division (no offspring) or a successful cell division (two daughter cells), that is *p*_2_ + *p*_A_ = 1. Given that *p*_2_ = 1 − *p*_A_, we can define the probability of survival *p*_A_ = Δ as the parameter of fitness for each single cell. In the case of a neutral branching process and at the initial state of the simulation (time *t*_2_) the probability of cell division is equal to the probability of cell death/differentiation for each single cell (Δ= 0.5).

We can translate this parameter Δ into a birth death process with *b*/*d* = *ω* using:

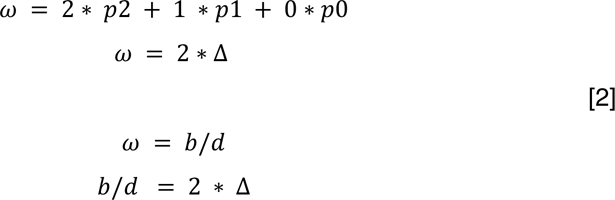

when Δ= 0.5 the birth death ratio *b*/*d* = 1.

The population grow exponentially when the probability of division (Δ) is 1. The probability of survival Δ used as a phenotype allow us to define a driver clone. A driver clone is a set of cells from the same evolutionary lineage which have the same Δ across them. This implies that they all have a shared set of mutations (very few or all) and the same survival probability. We also define an immunogenic clone, defined by the presence of an ancestral cell that acquired at least one immunogenic mutation.

#### 1.3 Cell genotypes and phenotypes

The genotype of each single cell is implemented as a vector storing the following information:

- Number of nonsynonymous mutations in driver, immune, escape, and passenger regions of the coding genome.
- Number of synonymous mutations in driver, immune and passenger regions of the coding genome.

At every successful cell division, each cell inherits the genotype from the parental cell, which is further modified by acquiring a new set of mutations. The number of new mutations is given by a Poisson distribution with mean *u* ∗ *L* with *L* = 50 ∗ 10^6^, *u* is the mutation rate per bp per cell division, and *L* is the length of the coding genome.

The phenotype of each single cell is implemented as a vector storing the following information:

- Fitness (probability of successful cell division) and strategy (passenger, driver, immunogenic or escape),

These phenotypes are the outcome of mutations present in the genotype vector. To estimate the target size and thus the vector of probabilities for passenger, driver, immunogenic, and escape mutations we used prior information and also explored different values. All tested values are described in supplementary table 1B.

#### 1.4 Cycle conditions

Our model requires the input of several parameters described in supplementary Table 1. We performed several simulations to account for the different phases described in figure 1. The parameters for the simulations are described in supplementary table 1B.

##### Mutation rate

Specifically, for MSS cases, we used a mutation rate per pb per cell division of 10^−8^. This value is a composite between the polymerase error and the DNA proofreading correction efficiency. For MSI and POLE cases we increased this value in one and two orders of magnitude respectively.

Initially, we estimated the probability of hitting a driver mutation (1%) based on the number of driver genes identified in a recent study using a pancancer dataset (~200 out of 20000 genes)^7^. We used 5% of the coding genome as immunogenic based on our recent analysis of immunogenic mutations^28^ and based on the length of all possible 9-mers defined as strong binders by NetMHCpan. The proportion of escape sites in the coding genome is unknown, thus we simulated different proportions ranging from 0.01% to 5%. In addition, we defined nonsynonymous mutations as: a) passenger mutations that do not have any effect on the phenotype, b) driver mutations increasing the probability of survival, b) immunogenic mutations that may elicit an immune response, d) escape mutations allowing the cell to hide from an immune attack. We assume that all synonymous mutations accumulate neutrally in the genome and define three types of synonymous a) synonymous mutations in neutral regions, b) in driver regions, and c) in the immunopeptidome. To simulate the dependency of the nonsynonymous to synonymous mutation ratio, dN/dS values, for global, driver and immune regions, we fixed the probability of synonymous mutations as 1/3 of the probability of the nonsynonymous mutations in the same locus. All these probabilities sum to one.

Then, each time a cell divides, each daughter cell inherits the parental genotype and an additional set of nonsynonymous and synonymous mutations based on the probability vector defined. Our model assumes infinite-sites and no-back mutation as used in previous studies^64^. Our model records the number of mutations for each mutation type, the probability vector for each of those mutations, the probability of survival and the probability of immune attack over time. We also store the parental relationship and we assign a new clone id only when the new genotype includes nonsynonymous driver different from the parental phenotype.

We stopped the simulation after 100 generations, consistent with the maximum number of cell divisions allowed by telomere shrinking, or when the population size reached a specific carrying capacity (2000 cells).

#### 1.4 Mutation effects

##### Phenotype 1 - Proliferation dynamics

We have developed a flexible framework to account for different models of fitness effects of driver mutations. We chose a model based on that to date we have mostly seen tumors having between 2-10 drivers, therefore at equilibrium we expect to reach an average of 5 clones each carrying a driver event or 1 clone carrying 5 driver events. Thus, we modelled the fitness increase by a driver event as a Gompertz function where driver events give different selective advantages based on the order of acquisition given by:

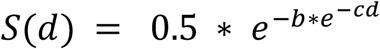

where *b* defines the displacement scale parameter of the Gompertz function, *c* defines the scale parameter on the fitness effect for each driver, and *d* is the number of driver events. We sample *b* from a normal distribution with mean 5 and *c* has a fixed value of 1.

Finally, at each time point each cell has a probability of survival defined by:

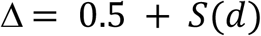

We must emphasize that the choice on these functions is perhaps one of the most important open questions in the field of cancer evolution. The fitness effects of combinations of multiple drivers (epistasis), the proportion of mutated sites leading to an increase/decrease of a selective advantage, and whether there is an upper boundary for the fitness increase remain largely unsolved and it was not the scope of this work. Here, we aimed to explore the effects of the immune system, the selective pressures and the emergence of escape mutations on a single unifying framework of tumor evolution.

##### Phenotype 2 - Immunoediting

To model the effect of the immune system during somatic evolution we assume two possible scenarios.

In the first, we allow cells to accumulate immunogenic mutations based on the size of the immunogenic genome and the mutation rate. Each immunogenic mutation will be detected by the immune system at an immune-mediated cell death rate of P_IS_ that will remove the immunogenic cell. This rate can be seen as the healthiness of the immune system or the capacity of T-cell recognition based on the diversity of the TCR repertoire (with 0 for immunosuppressed to 1 for immunocompetent, alternatively this can be seen as low recognition or high recognition potential by TCRs). By simplifying this value to an external probability independent of the genome, it allows us to model the effect of the microenvironment.

In the second, we define a function that at every generation calculates how many cells in a given clone are immunogenic (at least one neoantigen) and if this number is greater than a selected cut-off value (50 cells in our model), we kill all cells from that clone given a certain probability (defined previously as P_IS_). An immunogenic cell carries at least one immunogenic mutation and have not acquired an escape mutation. When a cell acquires an escape mutation, the immune system will no longer attack this cell.

#### 1.5 dN/dS computation

To estimate the dN/dS ratio we fixed the initial probabilities of occurrence of nonsynonymous mutations to be three times higher than the occurrence of synonymous mutations, as naturally observed in the coding portion of the human genome.

In general, in the first cellular divisions the number of synonymous mutations is close to 0 for many cells making the calculation of dN/dS implausible (infinite). We calculated the dN/dS for all mutations (global dN/dS), driver mutations (driver dN/dS) and immunogenic mutations (immune dN/dS) by adding up the observed counts in the alive cells at a given time t.

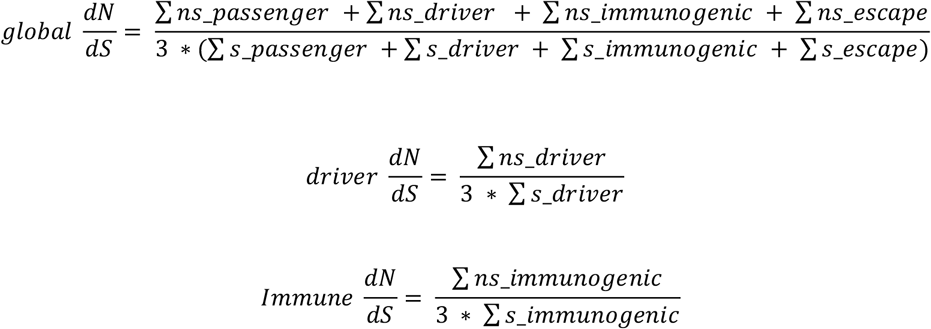

#### 1.6 Frequency dN/dS

To estimate the dN/dS ratio using a specific mutation frequency cut off we simulated sequencing by giving a mutation ID to each new mutation acquired during the stochastic branching process. We determine the cell-specific mutation by implementing an algorithm that walks along the lineage of a cell and concatenate all inherited mutations. We build a matrix of all alive cells and mutations at the last time point. We then were able to filter out variants present in less than any predefined threshold. For the driver section we used 0.01%, 1%, 2%, 3%, 4%, 5%, 10%, 25%, and 50% as frequency cut-offs. For the immune section we used 0.1%, 1%, 5%, 10% and 50% as frequency cut-offs. To estimate dN and dS we assigned each inherited mutation a unique id. Given that each mutation has two labels, a first label defined as 1) nonsynonymous and synonymous, and a second label defined as a 2) passenger, driver and immunogenic. This allowed us to calculate a global, driver, and immune dN/dS accordingly. Then, each simulation consisted of N number of cells with a specific number of nonsynonymous and synonymous driver, immunogenic, and passenger mutations.

#### 1.7 dN/dS confidence Intervals for frequency or cancer cell fraction cut-offs

When performing the analysis using frequency cut-offs, we pulled simulations together similar to what is done in cohort studies when all nonsynonymous and synonymous mutations are pulled together. To estimate the confidence interval for this analysis, we used the rateratio.test function from R package rateratio. This function calculates the p-value and the confidence interval for the rate of two Poisson ratios. It uses the uniformly most powerful ratio test available for R^65^.

### 2.0 TCGA Data

We first obtained somatic calls of TCGA data from GDC. This dataset consisted of 10202 samples across 33 tumors types. We then selected 19 tumor types tumor types that had been analysed in Rooney et al^51^ in order to compare our results of immune dN/dS to the immune cell scores. Rooney et al provided the per patient values of several normalized scores for immune cells. We calculated the median value for each score within each tumor type. The final list analysed consisted of 8543 samples across 19 tumor types. TCGA data was then re-annotated using ensembl-VEP release 89. COAD (Colon adenocarcinoma) and READ (Rectum Adenocarcinoma) were merged into CRC.

HLA-binding Mutation Ratios (HBMR) and simulated HBMRs were obtained from the supplementary material in Van Den Eynden et al^16^ available for 19 tumor types. Hartwig somatic calls and metadata were obtained from Hartwig Medical Foundation under license agreement DR-075.

#### 2.1 Selection On PRotein ANotated regiOns, SOPRANO

SOPRANO was developed on top of the method developed in Zapata et al 2018 to calculate selection in VEP annotated files and is freely available in github.com/luisgls/SOPRANO. It estimates dN/dS values in a target region (ON-target) and in the rest of the proteome (OFF-target) using a trinucleotide context correction (SSB192) or a 7-nucleotide context (SSB7). It allows the option to include or exclude cancer driver genes, as well as, randomizing the target region to calculate a background distribution of a matching size region. Given that it uses a set of Ensembl transcript identifiers and their respective FASTA file it allows calculation of dN/dS in any genome irrespective of the version. We ran SOPRANO on 33 tumor types and deposited the results for each tumor type in Synapse (syn22149238).

#### 2.2 Immunopeptidome and patient specific HLA

We downloaded a set of protein coding transcripts with HGNC symbol from Ensembl Biomart. We obtained all transcript lengths and run bedtools makewindows to get all possible overlapping 9-mers. We then obtained the FASTA sequence for each of all 9-mer and run netMHCpan4 using a list of HLA-alleles. This list of HLA-alleles was restricted to those that have more than 1% population frequency in a list of 1277 samples from the 1000K cohort. We selected all possible strong binders which had a mean and a median expression above 1FPKM. We obtained expression values for different tissues from the human protein atlas (downloaded on 05/10/2018).

### 3.0 Analysis of Metastatic cohort pre-immunotherapy from Hartwig Medical Foundation

We obtained the somatic mutation data from the Hartwig Medical Foundation cohort (HMF) under license agreement DR-075. The data we used for this manuscript consisted on 308 metastatic patients that underwent immunotherapy post-biopsy and that had recorded clinical response in “first response” column from the metadata. Mutation types that were not classified as synonymous, missense, start_lost, stop gained, stop lost or frameshift mutation were excluded. We removed indels and reannotated SNVs following our pipeline to obtain high confidence calls for a predefined set of ensemble transcripts (~20000 genes). We then rerun ensemble VEP using version 90 for Grch37 and parse the file using VATools V1.0.0. We uploaded the final annotated file used for the rest of the manuscript for each of the 308 patients to Synapse.

It is important to note that the raw clinical data was supplied by HMF and final consistency checks are still to be performed. The response evaluations were not performed as part of a clinical trial and the timing of the evaluations was variable. We classified patients into responders and non-responders based on the first response recorded after treatment was initiated. The group of responders consisted of those that were labelled complete response (CR, 1 case), or partial response (PR, 78 cases). Those that were labelled stable diseases (SD, 98 cases) or progressive disease (PD, 131 cases) were classified as non-responders. To keep consistency with other studies, there were 79 cases with no data, two cases classified as clinical progression, four cases classified as ND, and 3 cases classified as Non-CR/Non-PD which were not included in the analysis. The timing from biopsy to response was not included. There were no other further classifications performed.

Escape genes were selected based on the list of Antigen Processing and Presentation Machinery (hsa04612) download from KEGG. In addition, we included escape genes used in Rosenthal et al^34^. We then classified responders and non-responders into “escaped” if there was a missense or a truncating mutation in one of these escape genes and into “adapted” otherwise.

All statistical tests were performed using R statistical language. Statistical tests were performed using Wilcoxon rank-sum test for two distributions or Kruskal-Willis test when more than two distributions were present using the R package ggstatsplot.

We ran SSB192 (github.com/luisgls/SSB-dNdS) with default parameters to determine gene and global dN/dS values and SOPRANO (github.com/luisgls/SOPRANO) using the bed file provided in the package for HLA-A0201 using 192-base pair correction. We calculated dN/dS for driver genes using the list of 196 genes provided in Martincorena et al^7^. We calculated SSB-dNdS and immune dN/dS in the four group categories. For the TCGA patient specific SOPRANO analysis we used the 4-digit code HLA type for each gene (HLA-A, HLA-B, and HLA-C). We concatenated all regions predicted to bind to netMHCpan4.0 as strong binders in those genes that have a median expression of more than 1FPKM calculated across the 33 TCGA cancer types.

