## Supplementary Figures for "*dN/dS* dynamics quantify tumour immunogenicity and predict response to immunotherapy"

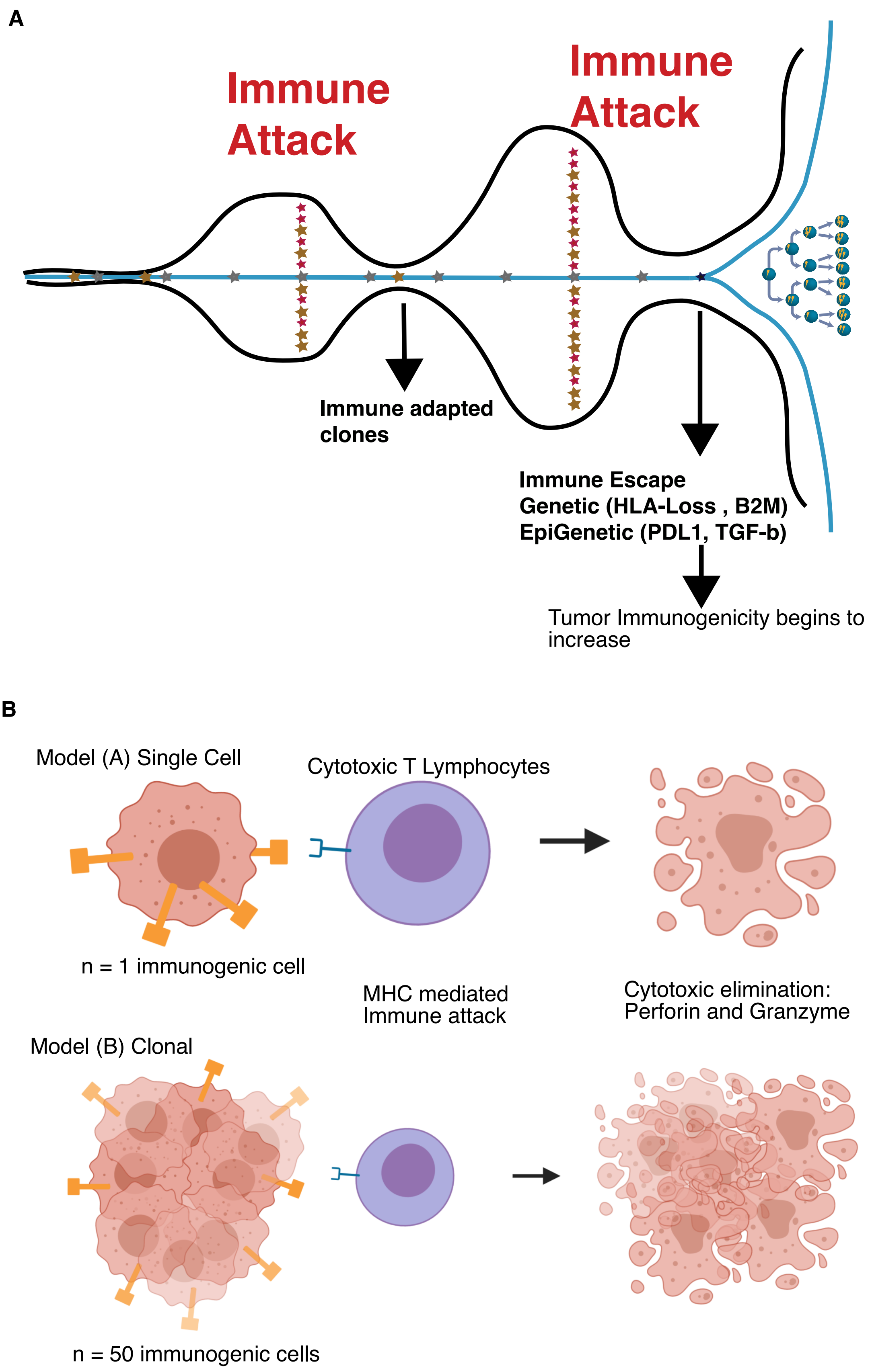
A
Immune
Attack
Immune
Attack
Immune adapted
clones
Immune Escape
Genetic (HLA-Loss , B2M)
EpiGenetic (PDL1, TGF-b)
Tumor Immunogenicity begins to
increase
B
Model (A) Single Cell
Cytotoxic T Lymphocytes
n = 1 immunogenic cell
MHC mediated
Immune attack
Cytotoxic elimination:
Perforin and Granzyme
Model (B) Clonal
n = 50 immunogenic cells

*Figure 1*

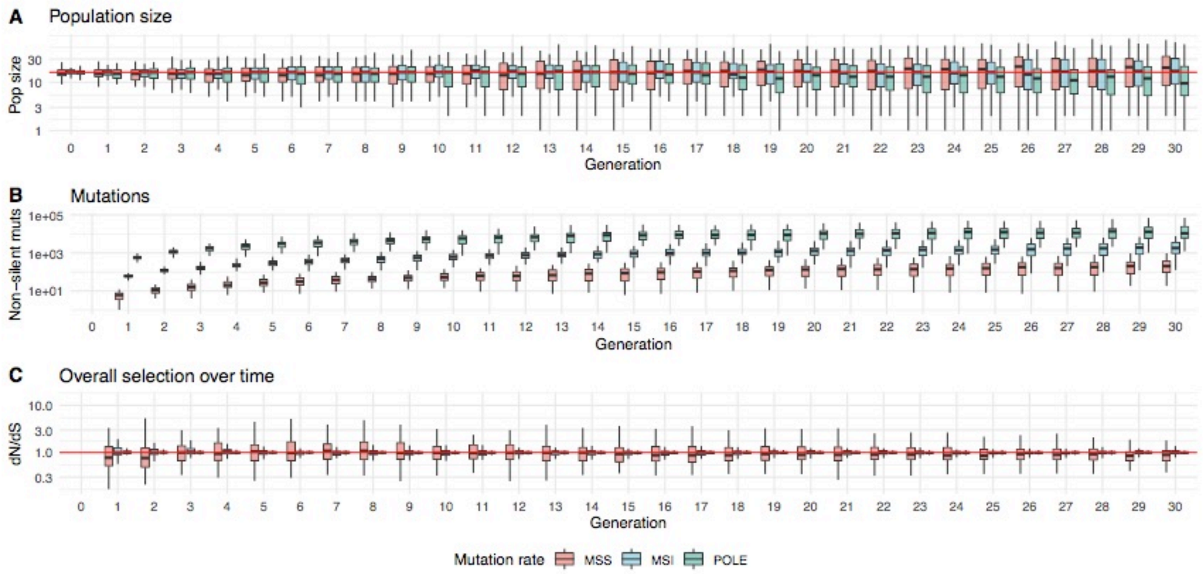
A Population size
Pop size
Generation
B Mutations
Non-silent muts
Generation
C Overall selection over time
dN/dS
Generation
Mutation rate MSS MSI POLE

*Figure 2*

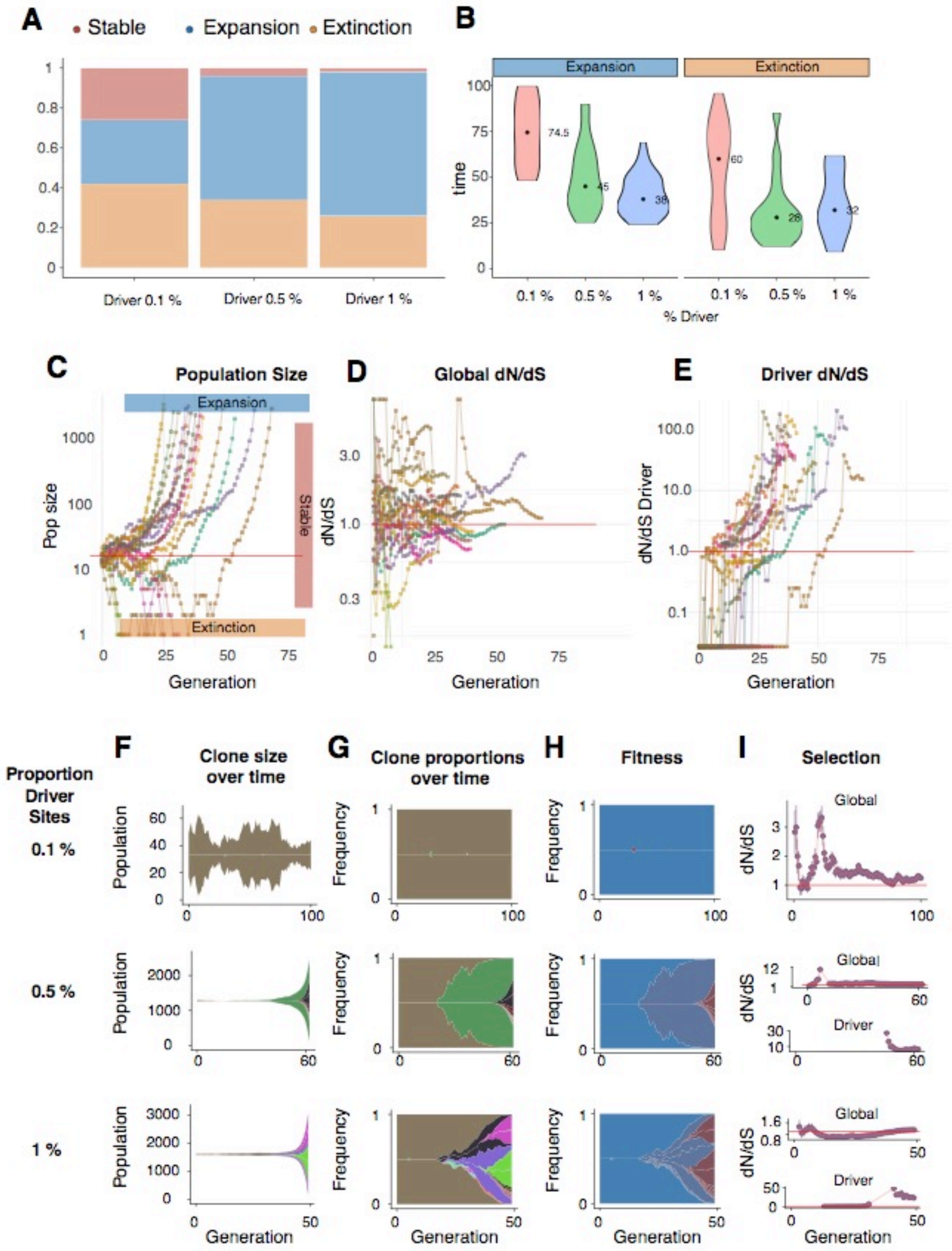
A
Stable
Expansion
Extinction
Driver 0.1 %
Driver 0.5 %
Driver 1 %
B
Expansion
Extinction
time
74.5
45
38
60
28
32
0.1 %
0.5 %
1 %
% Driver
C
Population Size
Expansion
Stable
Extinction
Pop size
Generation
D
Global dN/dS
dN/dS
Generation
E
Driver dN/dS
dN/dS Driver
Generation
Proportion Driver Sites
0.1 %
0.5 %
1 %
F
Clone size over time
Population
Generation
G
Clone proportions over time
Frequency
Generation
H
Fitness
Frequency
Generation
I
Selection
Global
Driver
dN/dS
Generation

*Figure 3*

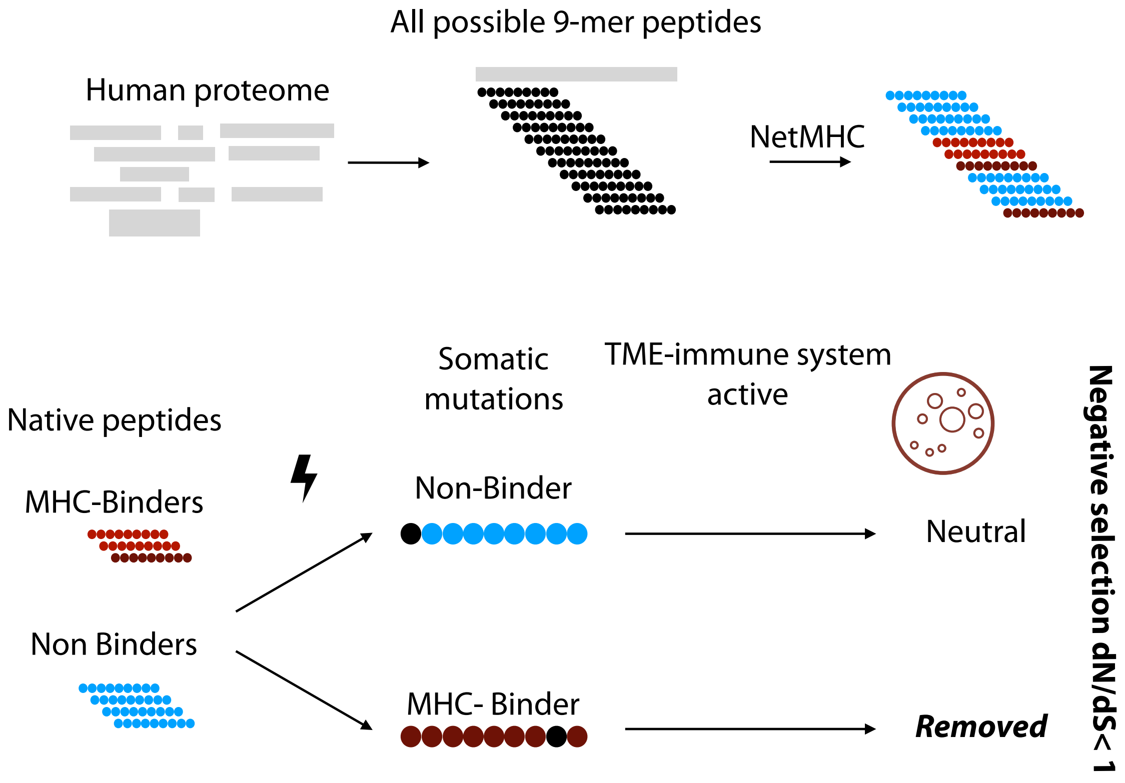
All possible 9-mer peptides
Human proteome
NetMHC
Somatic
mutations
TME-immune system
active
Native peptides
MHC-Binders
Non-Binder
Neutral
Non Binders
MHC- Binder
Removed
Negative selection dN/dS< 1

*Figure 4*

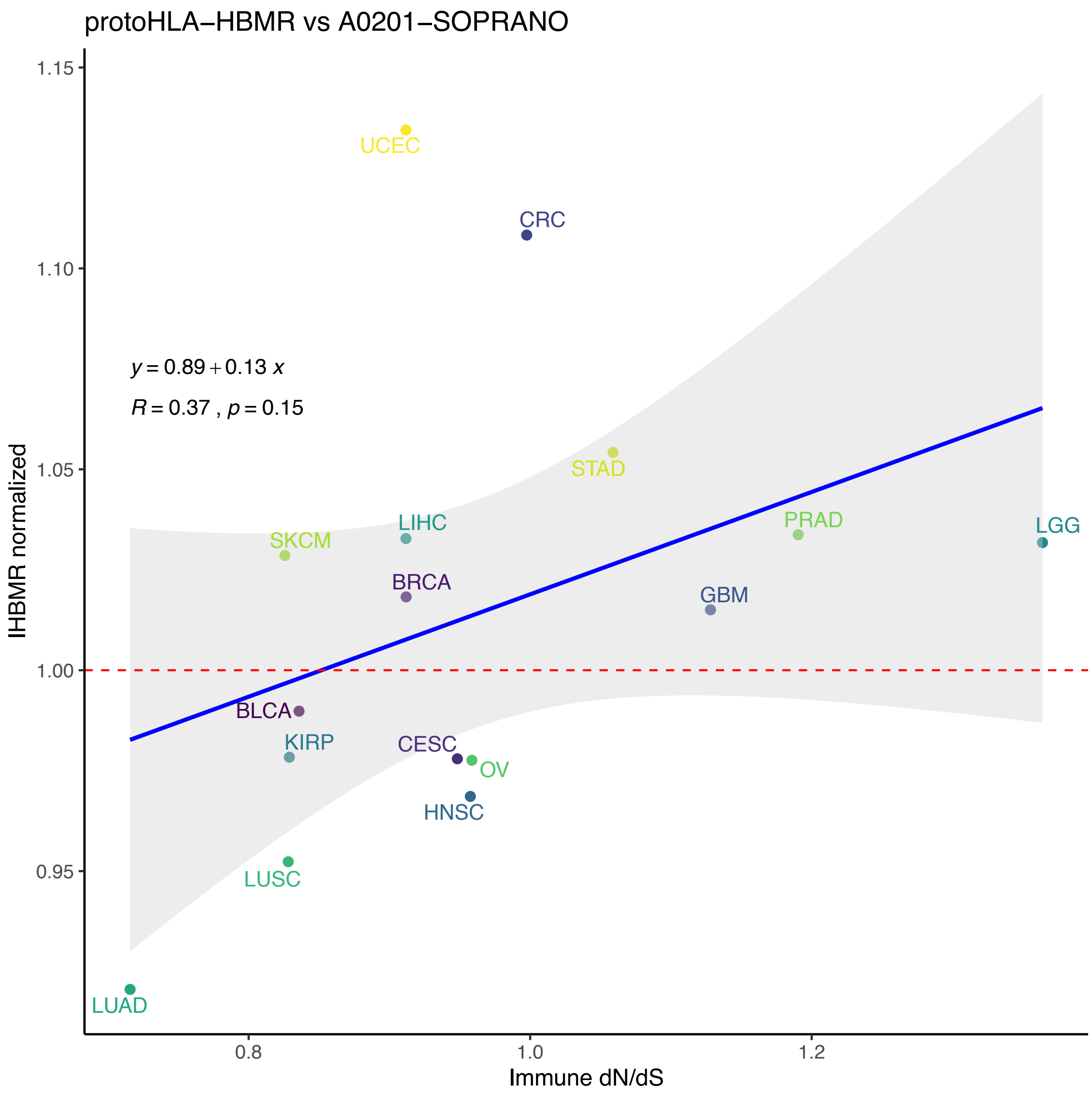
protoHLA−HBMR vs A0201−SOPRANO
1.15
1.10
1.05
1.00
0.95
IHBMR normalized
y = 0.89 + 0.13 x
R = 0.37 , p = 0.15
UCEC
CRC
STAD
LIHC
SKCM
PRAD
LGG
BRCA
GBM
BLCA
KIRP
CESC
OV
HNSC
LUSC
LUAD
0.8
1.0
1.2
Immune dN/dS

*Figure 5*

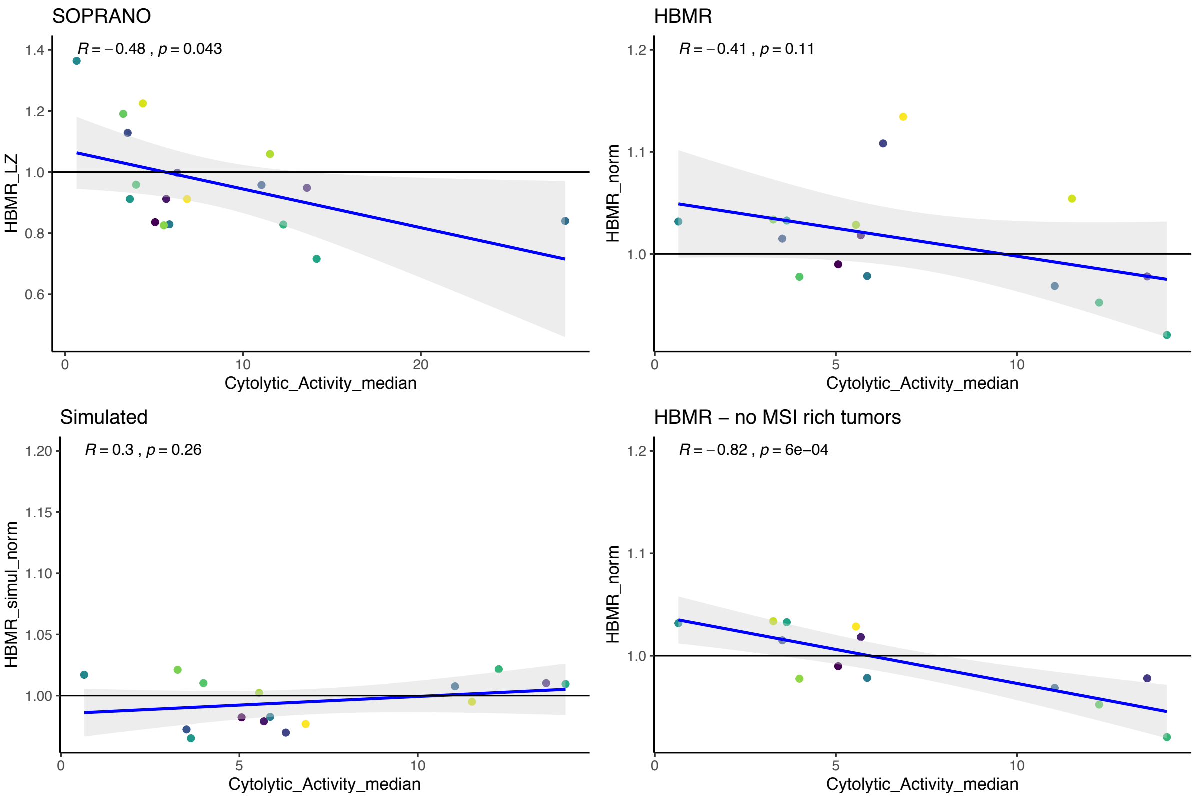
SOPRANO
R = −0.48 , p = 0.043
HBMR_LZ
Cytolytic_Activity_median
HBMR
R = −0.41 , p = 0.11
HBMR_norm
Cytolytic_Activity_median
Simulated
R = 0.3 , p = 0.26
HBMR_simul_norm
Cytolytic_Activity_median
HBMR − no MSI rich tumors
R = −0.82 , p = 6e−04
HBMR_norm
Cytolytic_Activity_median

*Figure 6*

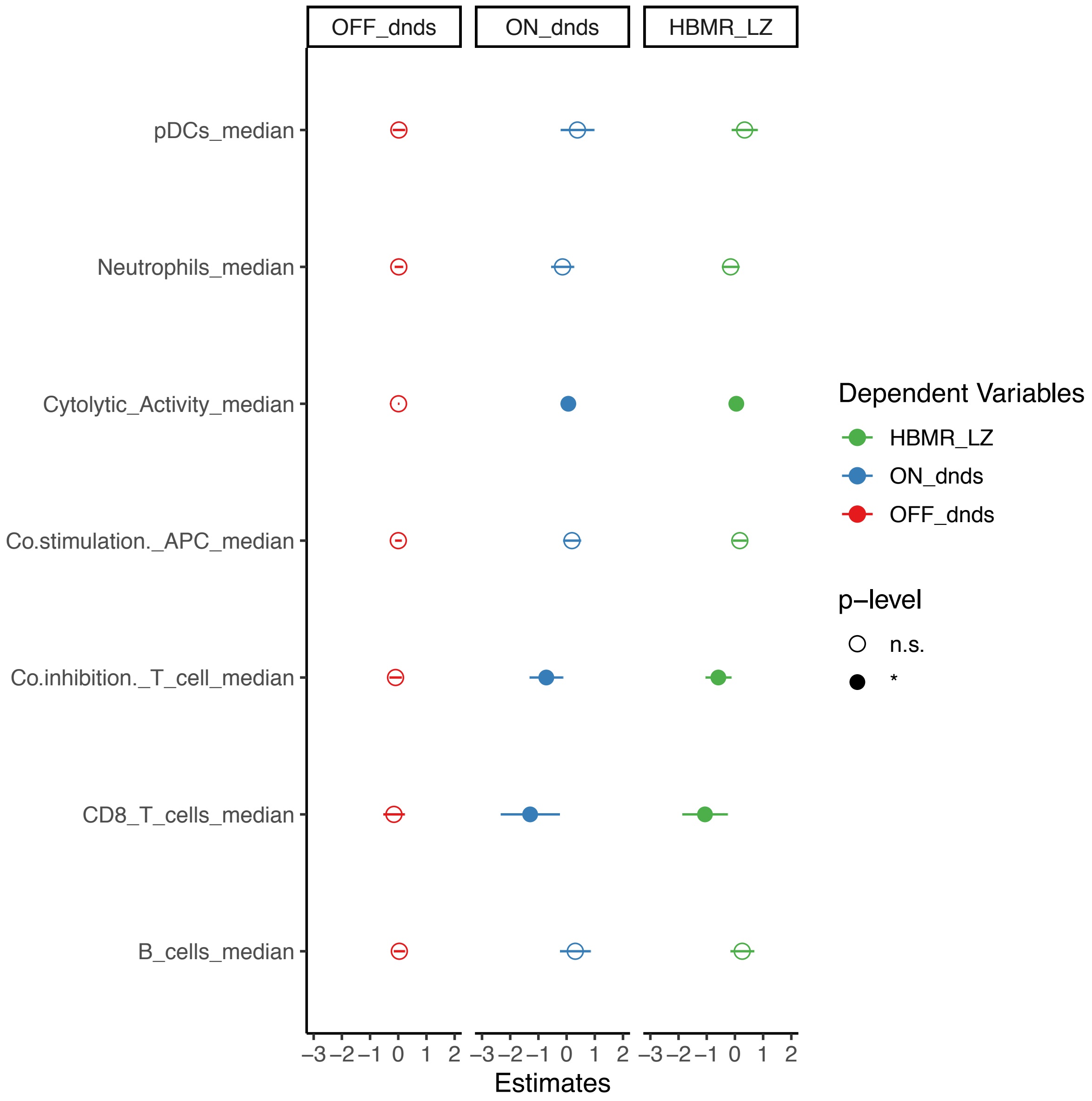

OFF_dnds
ON_dnds
HBMR_LZ
pDCs_median
Neutrophils_median
Cytolytic_Activity_median
Co.stimulation._APC_median
Co.inhibition._T_cell_median
CD8_T_cells_median
B_cells_median
−3 −2 −1 0 1 2
Estimates
Dependent Variables
HBMR_LZ
ON_dnds
OFF_dnds
p−level
n.s.
*


*Figure 7*

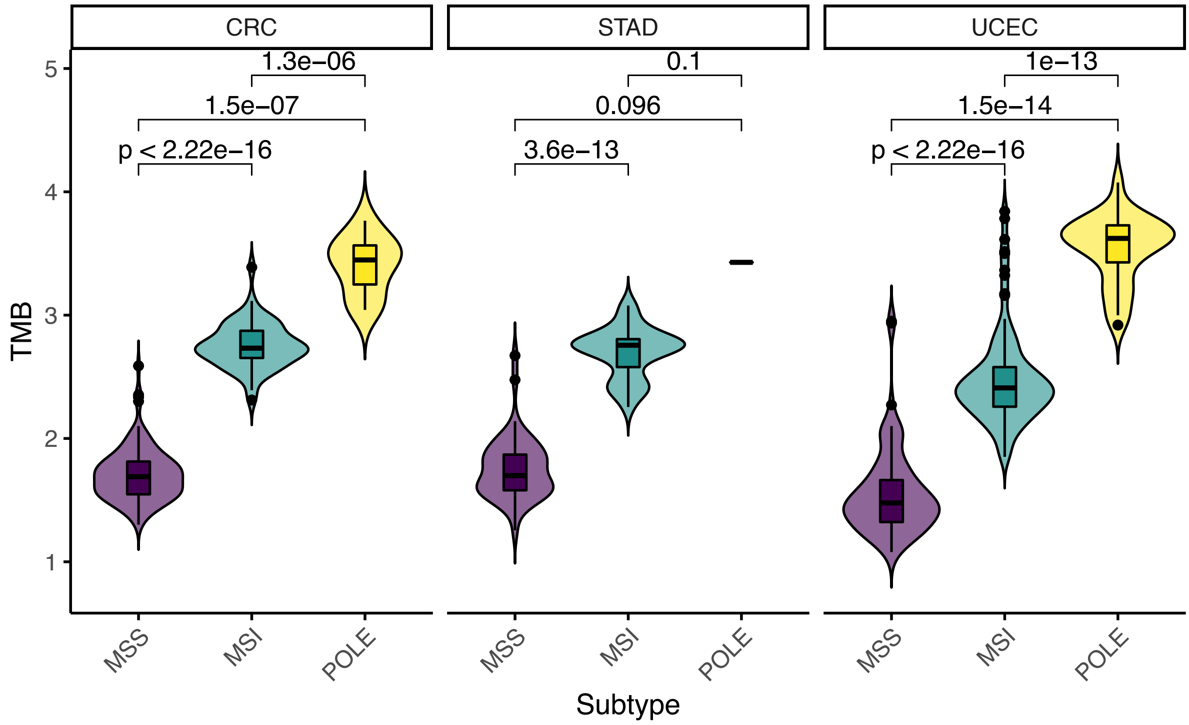


*Figure 8*

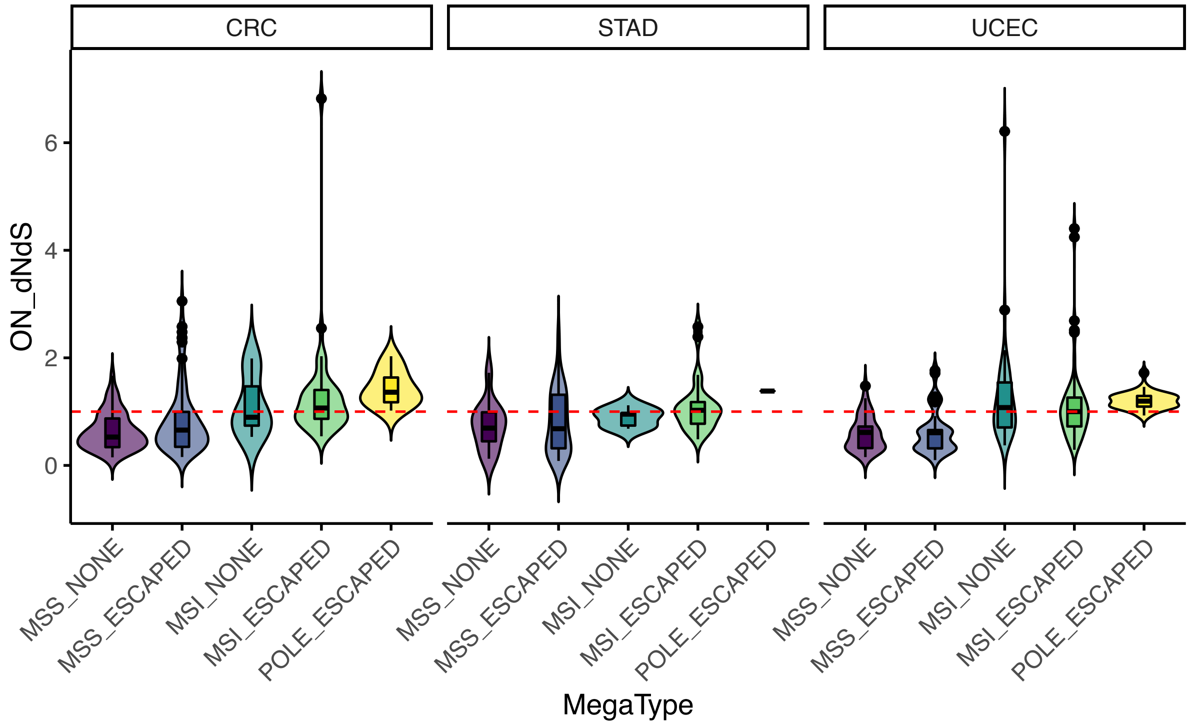


*Figure 9*

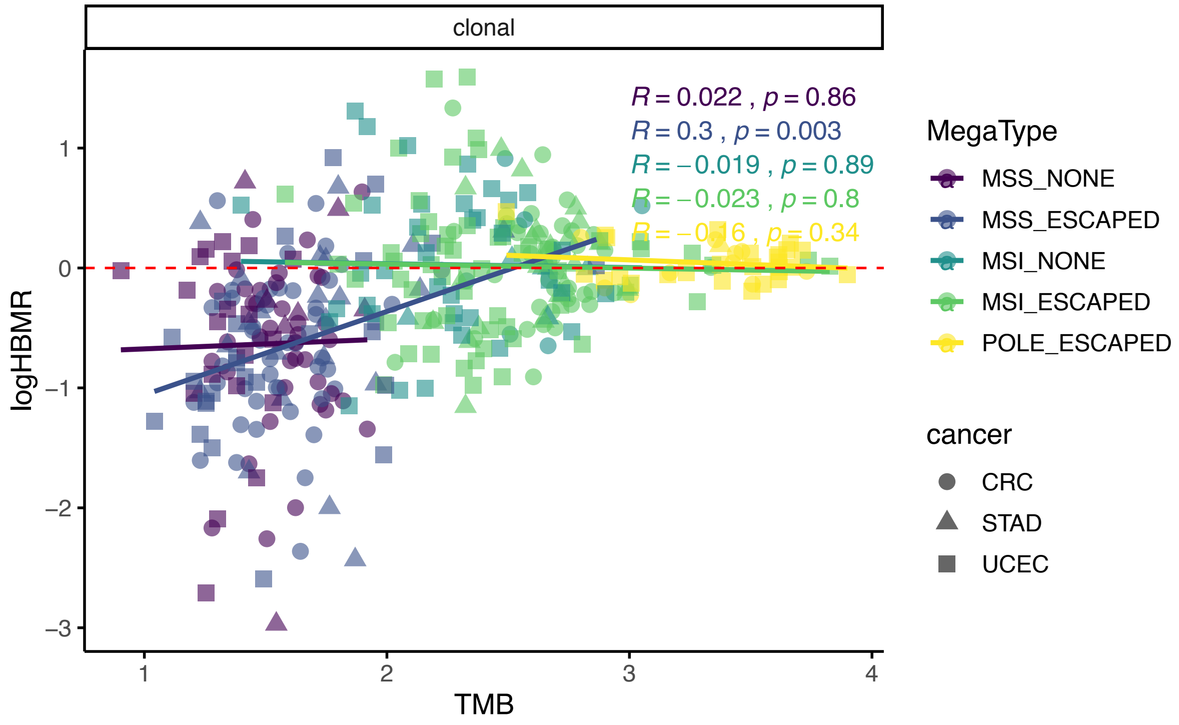


*Figure 10*

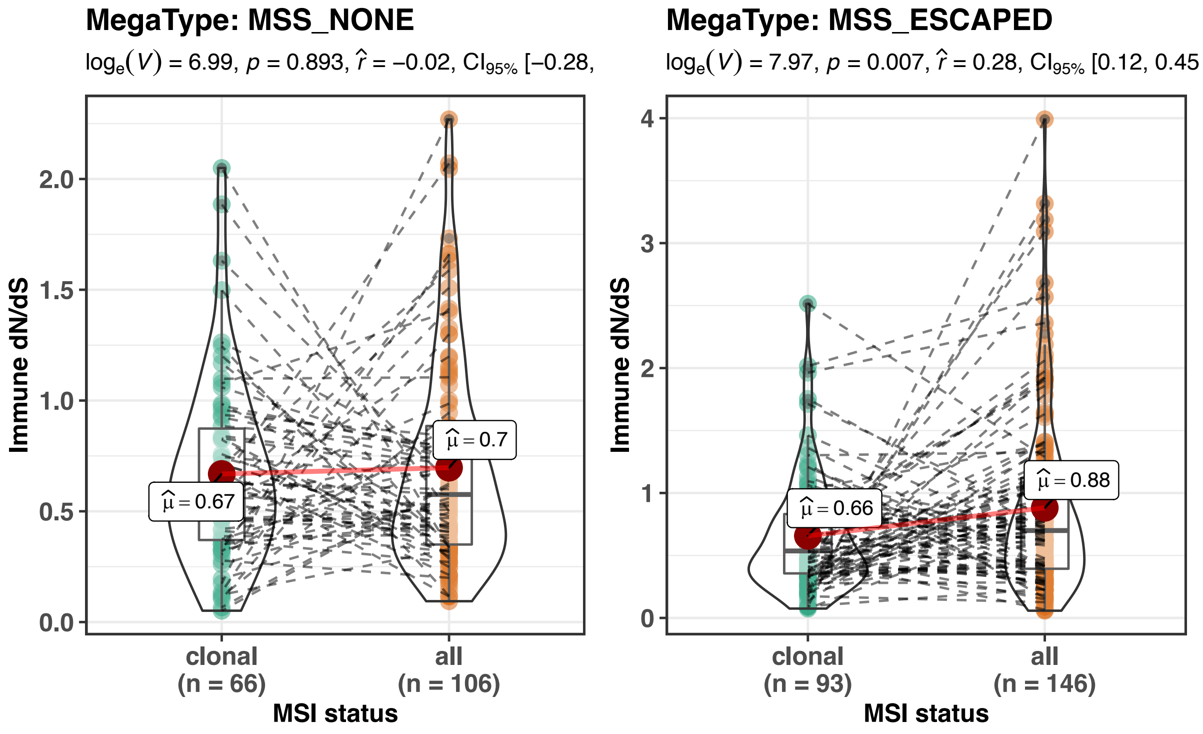


*Figure 11*

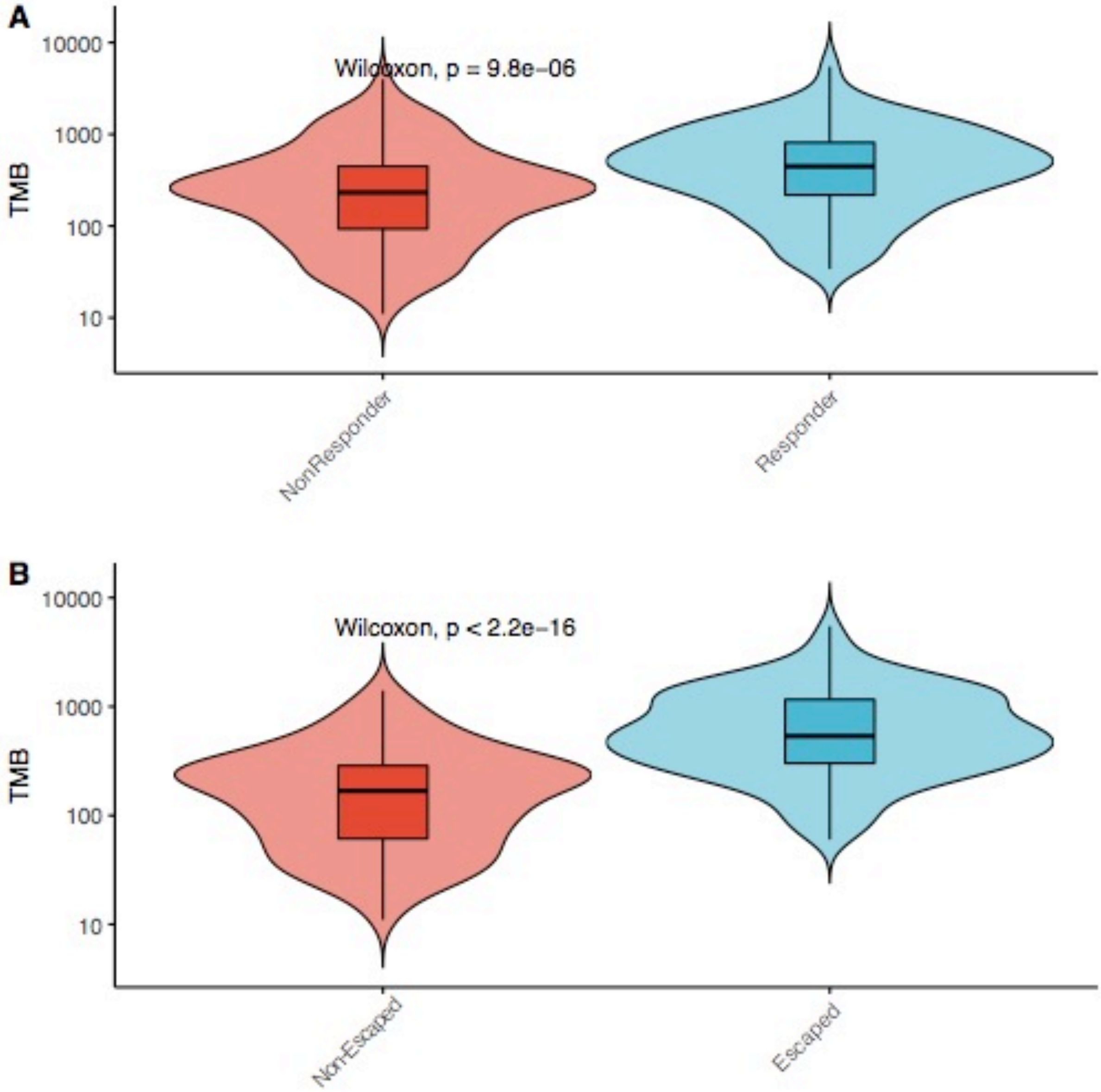

A
B
Wilcoxon, p = 9.8e−06
Wilcoxon, p < 2.2e−16
TMB
10000
1000
100
10
NonResponder
Responder
Non-Escaped
Escaped


*Figure 12*
